# TSWIFT, a novel method for iterative staining of embedded and mounted human brain sections

**DOI:** 10.1101/2023.09.19.558493

**Authors:** Corey M Porter, Matthias C. Truttmann

## Abstract

Comprehensive characterization of protein networks in mounted brain tissue represents a major challenge in brain and neurodegenerative disease research. In this study, we develop a simple staining method, called TSWIFT, to iteratively stain pre-mounted formalin fixed, paraffin embedded (FFPE) brain sections, thus enabling high-dimensional sample phenotyping. We show that TSWIFT conserves tissue architecture and allows for relabeling a single mounted FFPE sample more than 10 times, even after prolonged storage at 4 °C. Using TSWIFT, we profile the abundance and localization of the HSP70 family chaperones HSC70 (HSPA8) and BiP (HSPA5) in mounted human brain tissue. Our results establish TSWIFT as an efficient method to obtain integrated high-dimensional knowledge of cellular proteomes by analyzing mounted FFPE human brain tissue.

## Introduction

Mammalian tissue is comprised of a diverse mixture of cell types, each expressing its own characteristic set of proteins. Immunohistochemistry and immunofluorescence imaging are among the most common and appropriate tools to study this complex diversity in human tissue. However, these techniques are limited in scope by the number of proteins that can be probed (stained) at one time. The introduction of methods relying on antibody-dye conjugates (*1*), antibody-oligonucleotide conjugates (*2–4*), mild tissue stripping, optimized fluorophore usage, and/or horseradish peroxidase (HRP)-reactive fluorophores have increased the number of proteins that can be stained and visualized simultaneously. However, the reliance on unconventional antibody conjugates, equipment, and proprietary dyes prevents a broader application of these methods. The recent development of elaborate tissue delipidation and preservation techniques offers a convenient solution to most of these restrictions. One of these methods is SHIELD (**S**tabilization under **H**arsh conditions via **I**ntramolecular **E**poxide **L**inkages to prevent **D**egradation), a technique established to preserve free-floating tissue sections and tissue pieces (*5*). This process involves first creating strong intra- and intermolecular bonds with a flexible polyepoxide, which strengthens tissue architecture without interfering with protein epitopes or changing tissue morphology, allowing for repeated cycles of staining. Then, the tissue is delipidated, reducing background fluorescence. Following SHIELD treatment, a single tissue sample can be iteratively stained, imaged, destained, and re-stained using conventional antigen-specific primary antibodies and dye-conjugated secondary antibodies (*5*). SHIELD and related concepts have been successfully used to prepare large tissue samples (e.g., whole mouse brains), fixed and fixed-frozen cells in multi-well plates (*1, 6, 7*), and non-mounted frozen tissue sections. However, many archival samples provided by human tissue banks are formalin fixed, paraffin embedded (FFPE), and are often pre-cut and mounted onto glass slides. This makes these samples incompatible with currently available tissue delipidation and iterative staining methods.

In this study, we describe TSWIFT (**T**issue Staining **W**ith **I**terative **F**luorescent **T**agging), a novel tissue processing and staining method to iteratively stain mounted FFPE brain tissue on glass slides. TSWIFT preserves protein antigens in mounted tissues and integrates the signal amplification aspect of using primary and secondary antibodies. We show that TSWIFT permits more than 10 subsequent staining-destaining-restaining cycles of a single human brain section, even when stored up to 6 months between consecutive processing cycles. Using TSWIFT, we iteratively stain mounted FFPE brain sections from Alzheimer’s disease, Amyotrophic Lateral Sclerosis (ALS), and Huntington’s disease patients as well as healthy controls to determine the abundance of major HSP70 family chaperones in the neocortex. Taken as a whole, we establish TSWIFT as the first tissue processing and staining method optimized for mounted FFPE human brain tissue. This method allows to obtain high-dimensional representations of cellular proteomes in human brain sections at subcellular resolution, which may accelerate our understanding of pathophysiological processes in the human brain.

## Methods

### Tissue sections

FFPE tissue samples were provided by the University of Michigan Brain Bank. Tissues were fixed in 10% neutral buffered formalin, processed into paraffin blocks, and stored at room temperature. Samples were cut into 5µm thick sections onto Fisherbrand Tissue Path Superfrost Plus Gold slides and baked at 65^°^C for 1 hour.

### SHIELD FFPE Preparation

The SHIELD FFPE procedure was adapted from Park, et al 2018(*5*). Mounted FFPE brain sections were heated at 60^°^C for 10min before being deparaffinized in xylene, hydrated in 100%, 95%, then 70% ethanol followed by deionized water. Slides were then incubated in a 7:1 mix of SHIELD-ON buffer and SHIELD Epoxy solution from LifeCanvas Technologies Inc. Slides were incubated in this solution for 6 hours at 4^°^C with gentle shaking, then moved to room temperature for 24 hours with continued shaking. Tissue dilipidation was performed in SDS clearing solution (300mM sodium dodecyl sulfate, 10mM boric acid, 100mM sodium sulfate, brought to pH 9.0 with sodium hydroxide) at 37°C with shaking for 24 hours. Slides were then incubated in phosphate or tris-buffered saline with 0.2% Triton X-100 (PBSX/TBSX) overnight at 37°C to remove SDS.

### Iterative Immunostaining

Antigen retrieval was done in citrate buffer (10mM citric Acid, 0.05% Tween 20, pH 6.0) in a Rosewill steamer model RHST 15001 for 25 minutes. Slides were then washed in Tris Buffered Saline (TBS) once. Sections were blocked in 10% FBS, 10%BSA with 0.2% Triton X-100 in TBS for 1 hour at room temperature with shaking. Primary antibodies (see supplementary Table S1) were diluted in 5% BSA, 0.2% Triton X-100 in TBS, covered with a small piece of parafilm, and incubated overnight at 4°C. Slides were washed 3x in TBS before applying secondary antibodies. Secondary antibodies were diluted in the same buffer as primary antibodies, along with DAPI (4′,6-diamidino-2-phenylindole) at 50ug/ml and incubated on sections in the same manner as the primary antibodies for 1 hour at room temperature. To prevent autofluorescence from any lipofuscin remaining post lipid-clearing, 0.5% Sudan Black in 70% ethanol was applied for 10 min. Coverslips were applied to tissue sections using ProLong Diamond Antifade Mountant (Invitrogen) and allowed to cure overnight before imaging. After imaging, coverslips were removed by incubating slides in 70% ethanol at room temperature or TBS at 37°C before destaining in destaining buffer as described in Park, *et al.,* 2018 (7mM SDS, 0.5M Tris HCl, pH6.0, 8 1% 2-mercaptoethanol) at 37 °C for 18 hours (*5*). Slides were washed three times 30 minutes in TBS before beginning the next round of staining, starting with the blocking step.

### Image acquisition and analysis

Stained slides were imaged using a Keyence BZ-X700 Microscope equipped with a CFI Nikon plain Apo 20x objective. Exposure times for each channel were set using the auto-exposure function in the BZ-X software and kept constant for the entire imaging cycle. Image analysis was done using BZ-X Analyzer and BZ-X Wide image Viewer from Keyence. Cyclic stains were analyzed by comparing small areas of the tissue in subsequent stains to first determine the viability of our approach. Once the technique was validated, small areas of each cycle of staining on the same tissue were compared to determine which markers co-localize. Stains of each chaperone of interest in various areas of the brain were analyzed for location within the area and for subcellular localization. To facilitate visual inspection of imaging results, color levels were automatically enhanced using the BZ-X Analyzer software, if necessary.

## Results

### TSWIFT allows for repeated staining and destaining with signal pattern retention

To establish an easily adaptable technique for iterative staining of mounted FFPE sections, we developed a new tissue clearance and staining protocol, incorporating commercially available SHIELD reagents (figure 1) (*5*). We first deparaffinized and rehydrated 5µm FFPE brain sections before epoxidation using SHIELD ON and SHIELD Epoxy buffers to strengthen tissue sections to withstand consecutive staining cycles. Next, we delipidated sections for 24 hours to reduce autofluorescence. Following a washing cycle, we then followed a traditional immunofluorescence staining protocol using Sudan Black (to reduce background fluorescence; see supp. figure 1), unconjugated primary antibodies, and fluorophore conjugated secondary antibodies.

**Figure 1.**
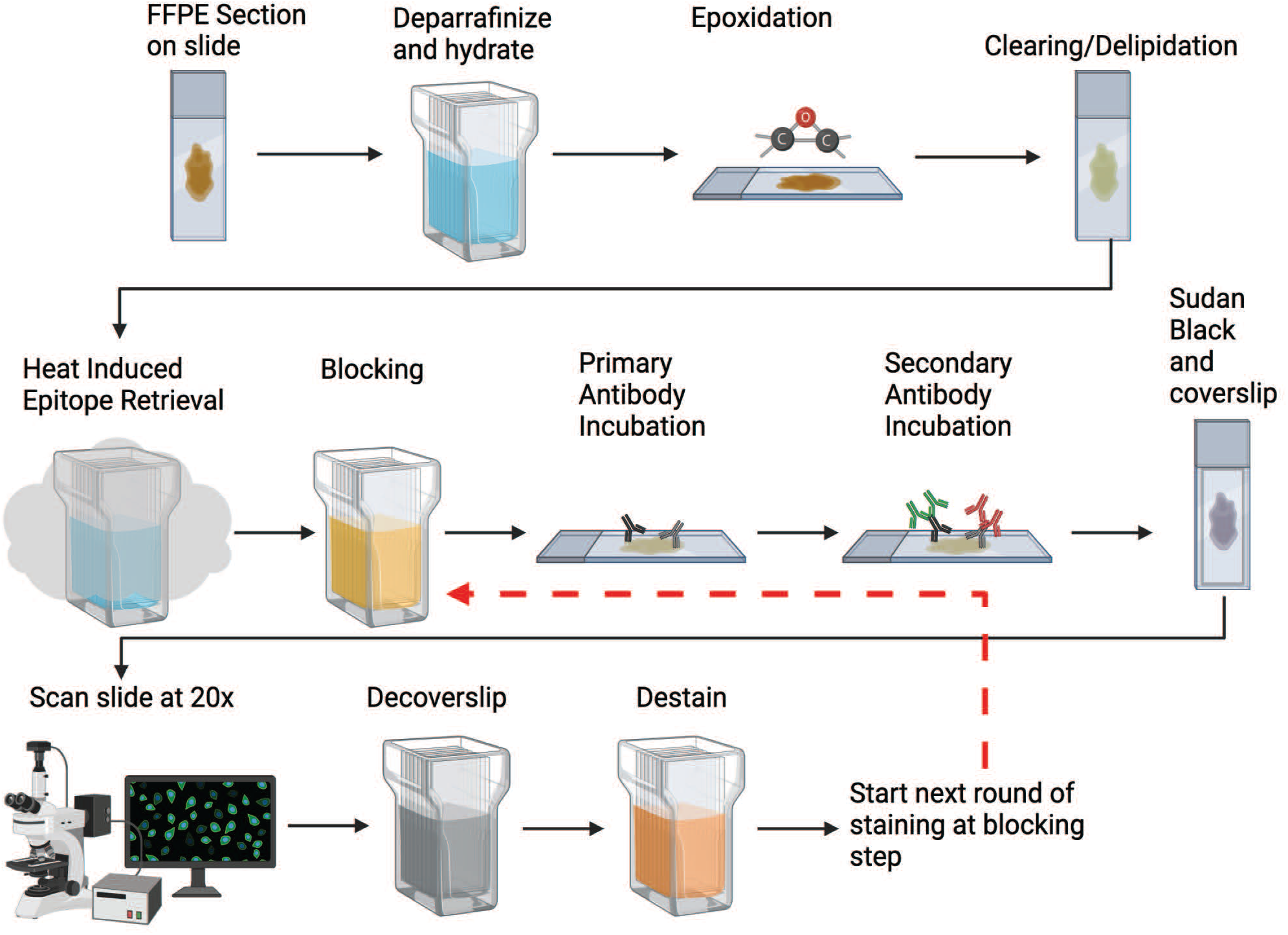
Schematic overview. A schematic representation of the TSWIFT protocol. Deparrafinization and rehydration are followed by epoxidation using SHIELD ON and Epoxy buffers to strengthen the tissue. Sections are then delipidated to reduce autofluorescence, followed by washing and a traditional immunofluorescence protocol including Sudan Black. The entire section is scanned at 20x before decoverslipping, destaining, and restarting the staining process at the blocking step.

To validate our new TSWIFT protocol on mounted FFPE sections, we conducted a series of experiments in which we processed and iteratively imaged, destained, imaged, and restained a cortex FFPE section, with the same primary and secondary antibodies (DAKO anti-GFAP; secondary fluorophore 488, Santa Cruz anti-Tau; secondary fluorophore 568) (figure 2, supp. figure 2-8). We found that even after 10 staining-destaining cycles, epitope patterns were well retained (supp. figures 2-8). Control imaging after each destaining step confirmed that little to no signal was retained (figure 3). Staining with only secondary antibodies in the fifth cycle and again using primary antibodies paired with secondary antibodies coupled to Alexa488 in the tenth staining cycle further confirmed efficient removal of antibodies during the destaining process (supp figure 4 and 6, figure 4). After destaining after cycle 10, we stored the test slide for 1 month at 4°C in TBS supplemented with 0.02% sodium azide. After this storage period, we stained and imaged the sample and found that epitope signal patterns were still intact (figure 5a-c). Upon destaining in cycle 13, we stored the sample for 4 months at 4°C. Staining and imaging of the sample following this 4-month storage period still resulted in consistent epitope patterns (figure 5d). Because of repeated coverslip removals, we observed minor sample disruptions (i.e., tears in the tissue section). Testing several cover slip removal methods, we found that removing the coverslip by soaking the slide in a PBS bath at 37°C with gentle shaking resulted in minimal tissue integrity loss. Together, these results establish that TSWIFT is an effective approach to iteratively stain mounted FFPE tissue sections with minimal signal loss and tissue disruption.

**Figure 2.**
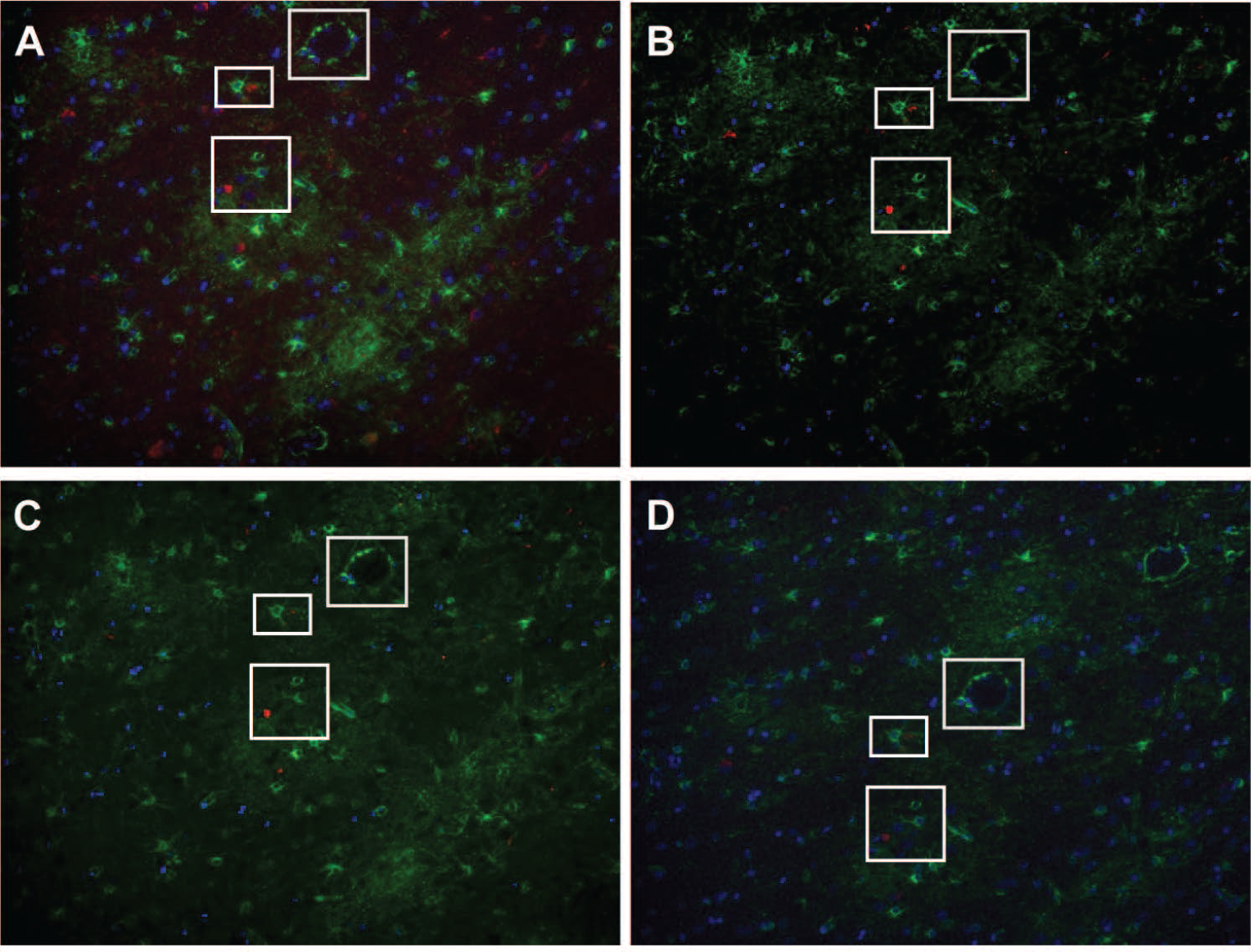
SHIELD protects tissue and allows for repeated staining of tissue with maintained signal. 20x images from cycles of staining post-SHIELD. All are stained with DAPI at 50µg/mL, Santa Cruz sc-58860 mouse mAb Tau (secondary 568) and DAKO Z0334 rabbit pAb GFAP (secondary 488). A: cycle 1, B: cycle 4, C: cycle 7, D: cycle 9.

**Figure 3.**
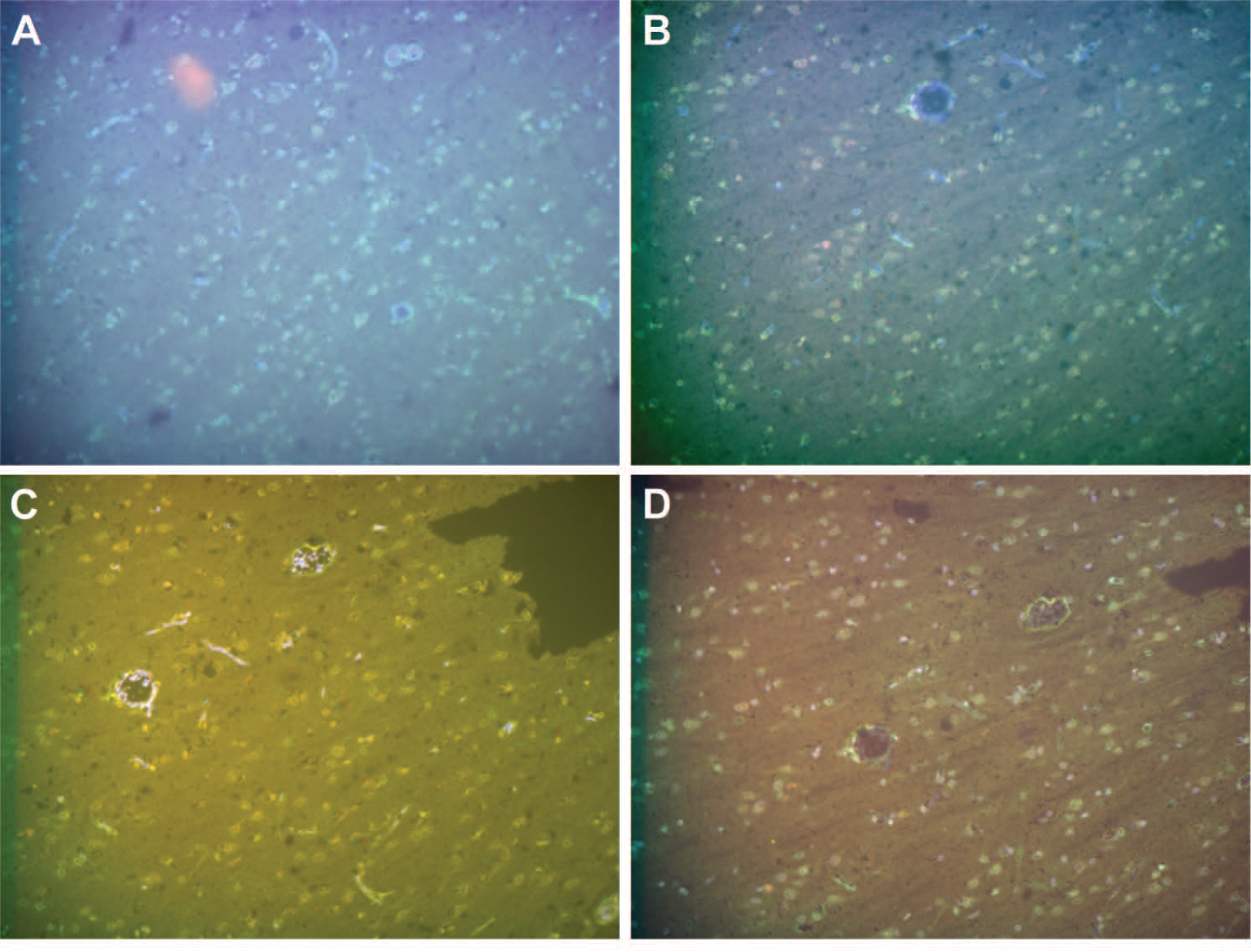
Destained sections show little to no retention of staining. Images taken at very long exposure times (up to 3.5s) confirm efficient tissue destaining and minimal tissue degradation. 20x images from destain cycles of staining post-SHIELD; A: cycle 1 destain, B: cycle 4 destain, C: cycle7 destain, D: cycle 9 destain.

**Figure 4.**
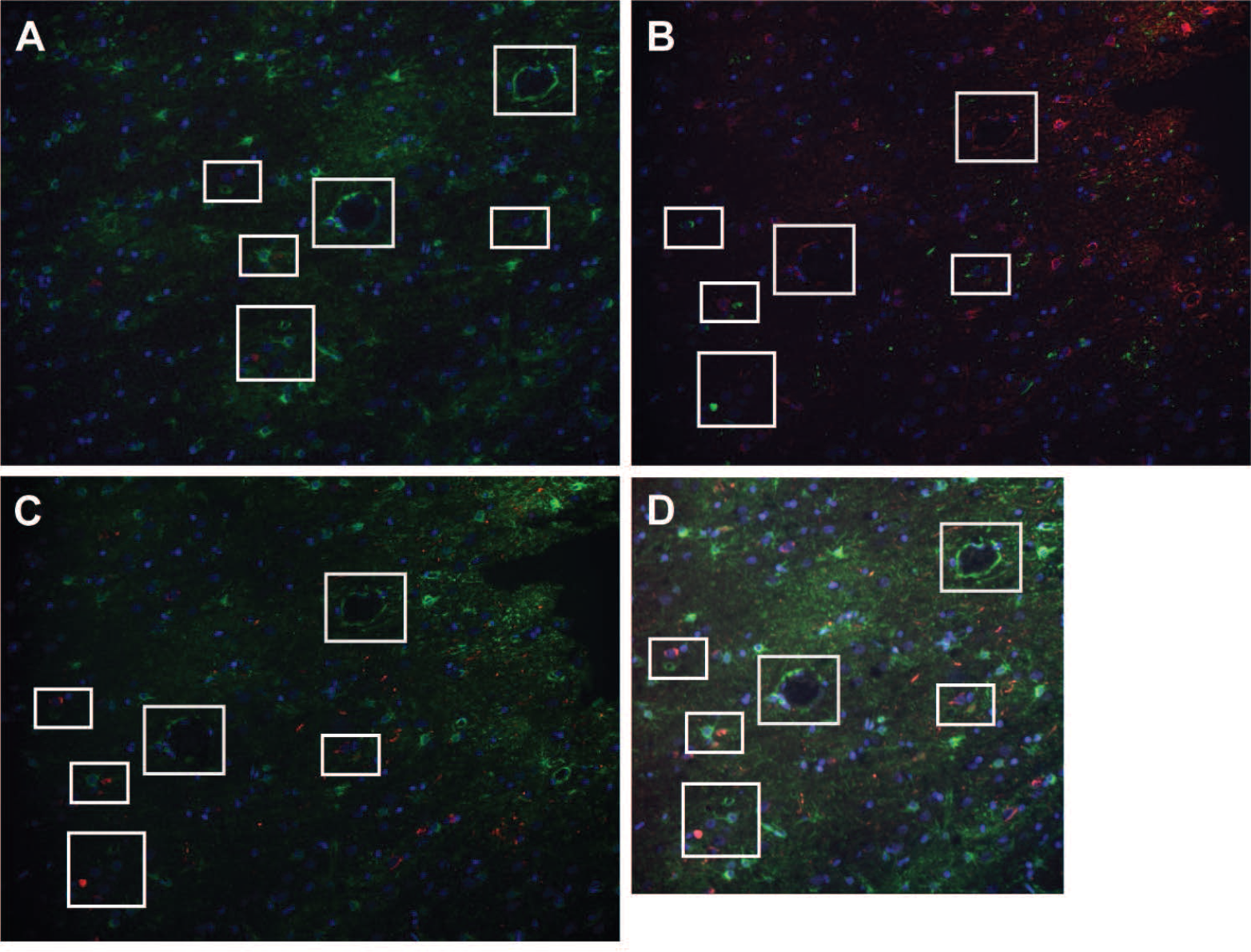
Residual secondary antibody deposits are not contributing to signal pattern. Images from cycle 9 and 10 of proof of concept staining. All are stained with DAPI, Santa Cruz sc-58860 mouse mAb Tau and DAKO Z0334 rabbit pAb GFAP. A: cycle 9 has Tau in red (568) and GFAP in green (488). B: cycle 10 has Tau in green (488) and GFAP in red (568). C: cycle 10 re-colored so Tau is red and GFAP is green D: cycle 10 re-colored so Tau is red and GFAP is green, and overlayed with cycle 9 (only area of overlap shown).

**Figure 5.**
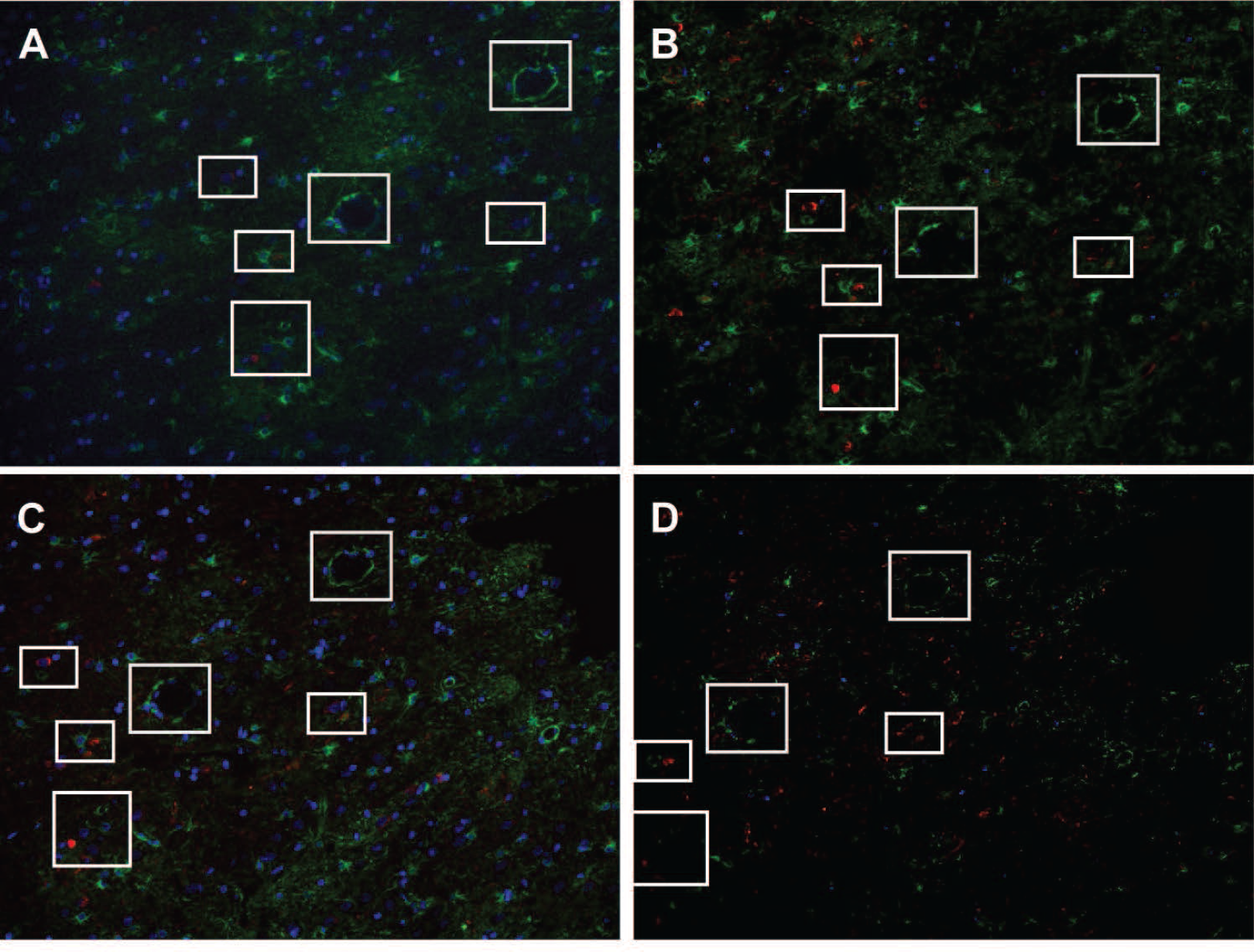
Sample storage at 4 °C does not impair processing. 1 month storage at 4 °C does not decrease staining, while 4-month storage decreases signal but maintains staining pattern. All are stained with DAPI, Santa Cruz sc-58860 mouse mAb Tau (secondary 568) and DAKO Z0334 rabbit pAb GFAP (secondary 488). A: cycle 9 (before 4 °C storage) B: cycle 11 (after 1 month 4 °C storage) C: cycle 13 D: cycle 14 after 4 months at 4 °C.

### Using TSWIFT to determine chaperone localization in human brain tissue

In a next step, we used TSWIFT to determine the distribution of the cytosolic HSP70 family chaperone, HSPA8 (HSC70) and the ER-resident HSP70 chaperone, HSPA5 (BiP), in brain sections from Alzheimer’s disease (AD), Huntington’s disease (HD), Amyloid Lateral Sclerosis (ALS), and control patients. We found that both HSPA8, and to a lesser extend HSPA5, are enriched in the grey matter of the cortex of all samples analyzed (HSPA8: 1 AD, 2 HD, 1 ALS, and 2 control; HSPA5: 1 AD, 1 HD, 3 ALS, 2 control) (figures 6-13). In the hippocampus, both HSPA8 and HSPA5 are enriched in cells populating the dentate gyrus (HSPA8: 2 AD, 2 HD, 2 ALS, 2 control; HSPA5: 1 HD) (figures 14-18). These results confirm and extend on previous work defining HSP70 family chaperone patterns in mammalian brains (*8, 9*).

**Figure 6.**
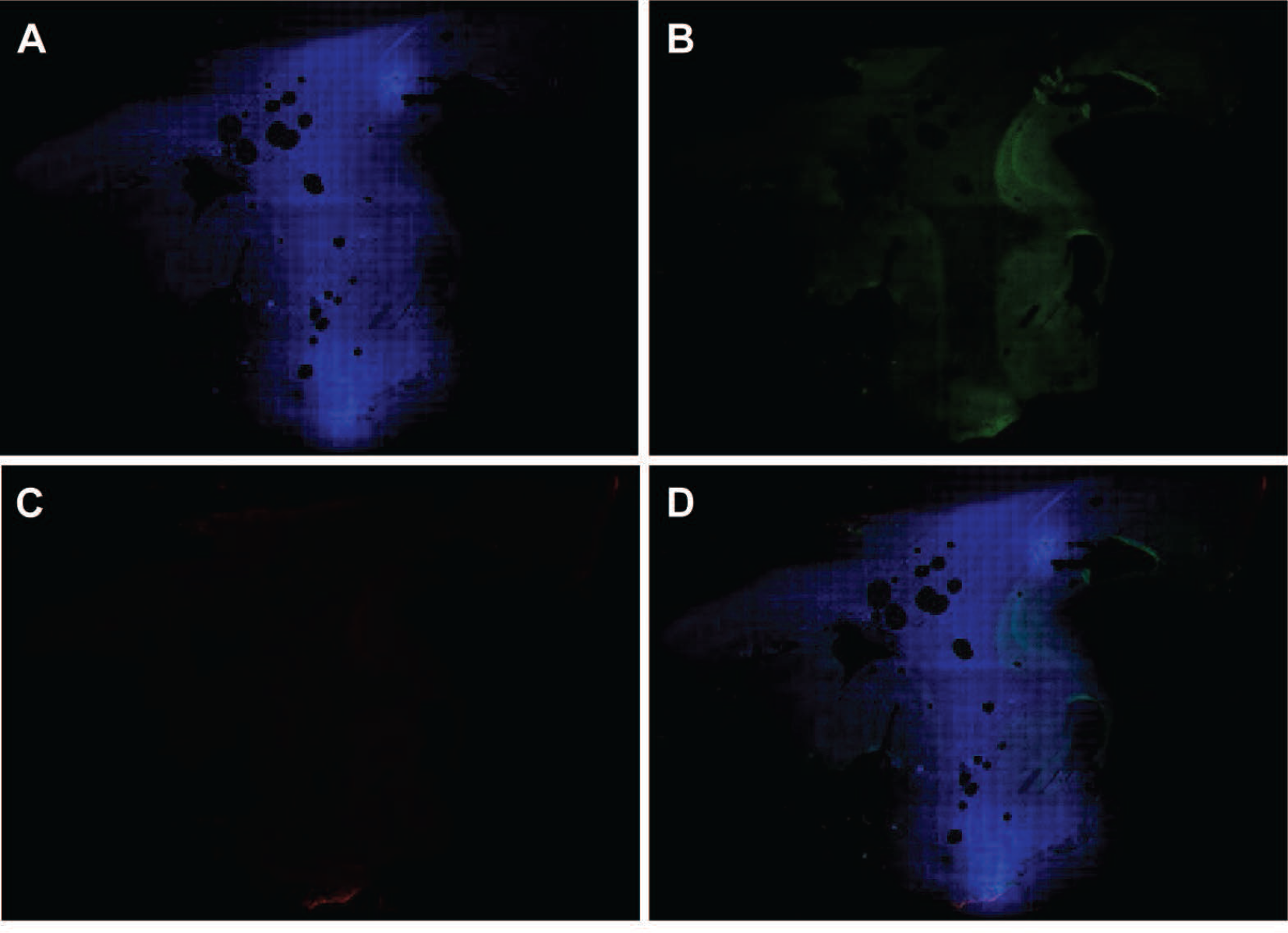
HSPA8 is more prevalent in the grey matter of a cortex with Alzheimer’s disease. Single channel and overlay images from whole slide scan of a cortex FFPE sample from an individual with Braak VI stage Alzheimer’s disease stained with DAPI, Santa Cruz 7298 mouse mAb HSPA8 (secondary 488) and ProteinTech 26975-1-AP rabbit pAb NeuN (secondary 568) antibodies. A: DAPI channel, B: 488 channel, C: 568 channel, D: overlay.

**Figure 7.**
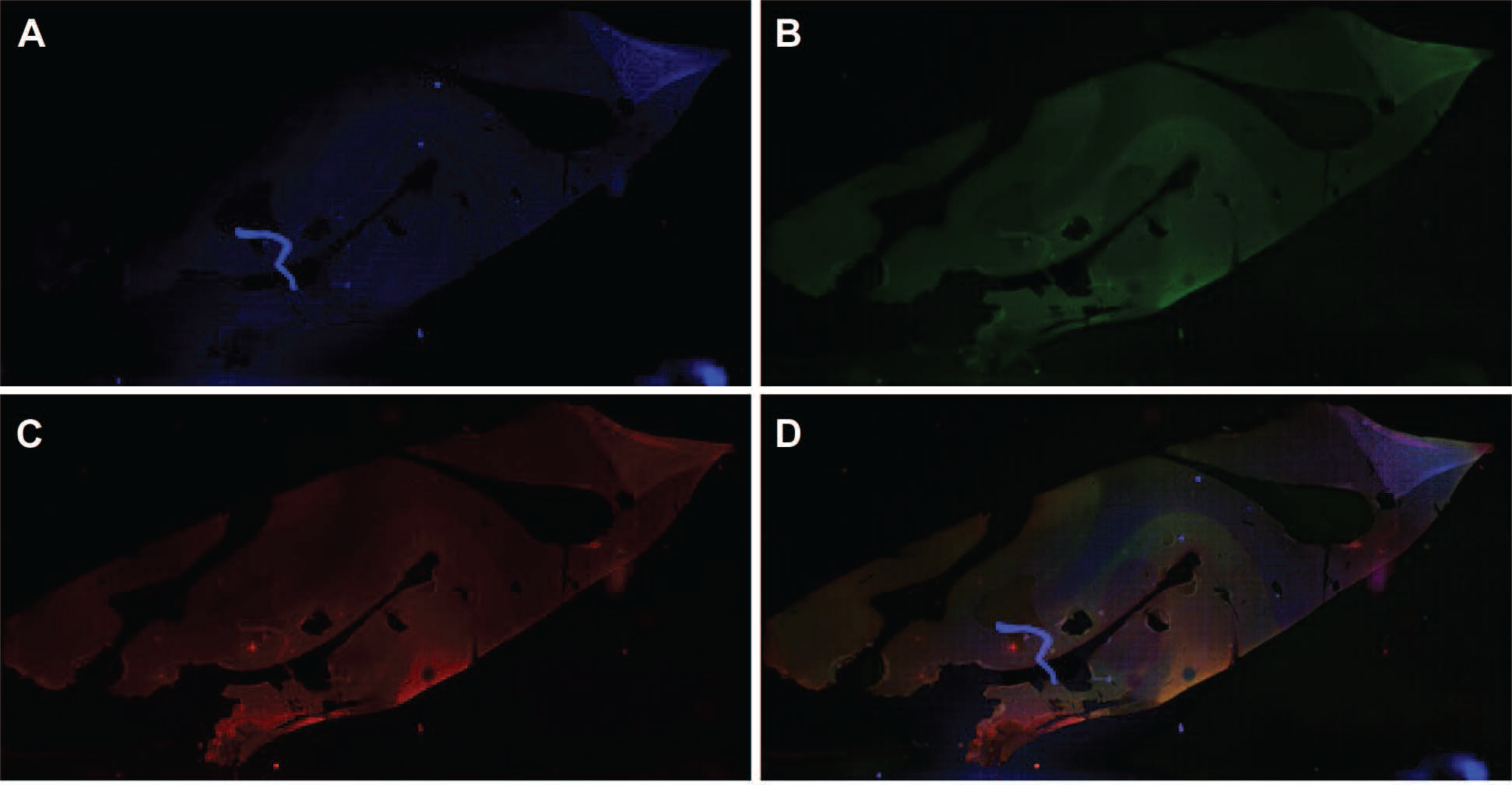
HSPA8 is more prevalent in the grey matter of a cortex from a Huntington’s disease case. Single channel and overlay images from whole slide scan of a segment of cortex from an individual with Huntington’s disease stained with DAPI, Santa Cruz 7298 mouse mAb HSPA8 (secondary 568) and Dako A0024 Rabbit pAb Pan human tau (multiple isoforms) (secondary 488) antibodies. A: DAPI channel, B: 488 channel, C: 568 channel, D: overlay.

**Figure 8.**
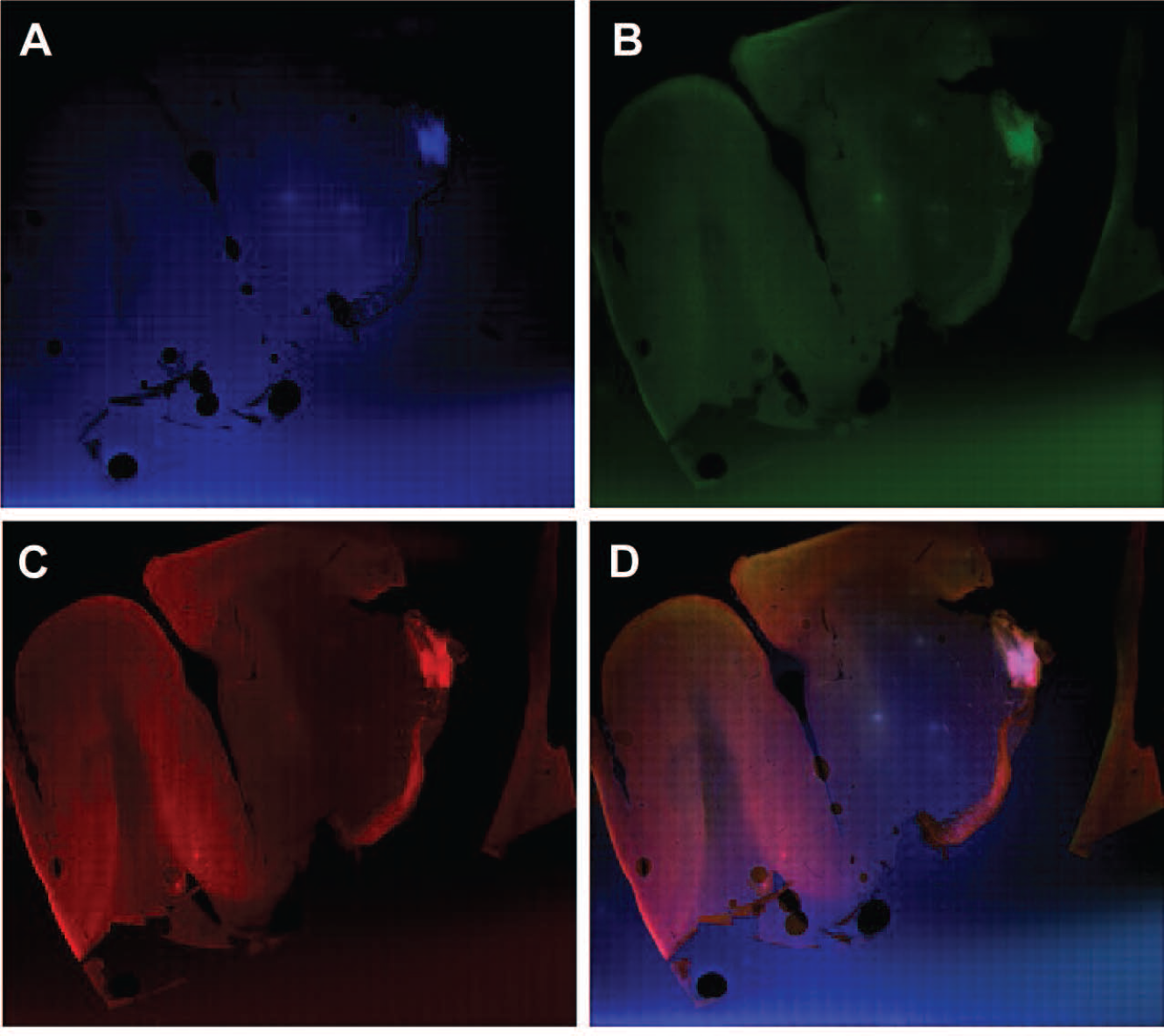
HSPA8 is more prevalent in the grey matter of a cortex with ALS. Single channel and overlay images from whole slide scan of a segment of cortex from an individual with Amyotrophic Lateral Sclerosis stained with DAPI, Santa Cruz 7298 mouse mAb HSPA8 (secondary 568) and Dako A0024 Rabbit pAb Pan human tau (multiple isoforms) (secondary 488) antibodies. A: DAPI channel, B: 488 channel, C: 568 channel, D: overlay.

**Figure 9:**
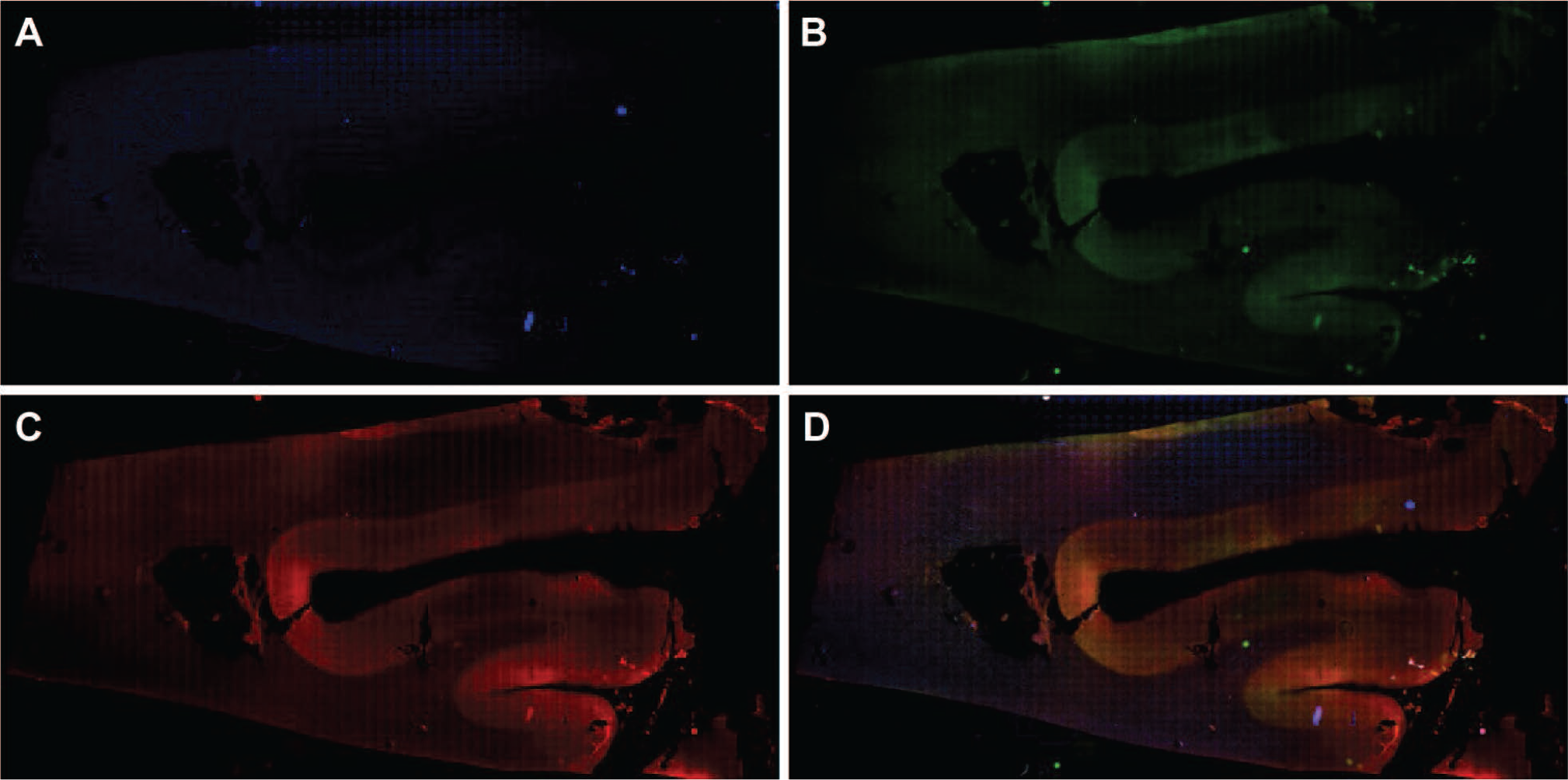
HSPA8 is more prevalent in the grey matter of a cortex from a donor with no neurological diagnosis. Single channel and overlay images from whole slide scan of a segment of cortex from an individual with no neurological diagnosis stained with DAPI, Santa Cruz 7298 mouse mAb HSPA8 (secondary 568) and Dako A0024 Rabbit pAb Pan human tau (multiple isoforms) (secondary 488) antibodies. A: DAPI channel, B: 488 channel, C: 568 channel, D: overlay.

**Figure 10.**
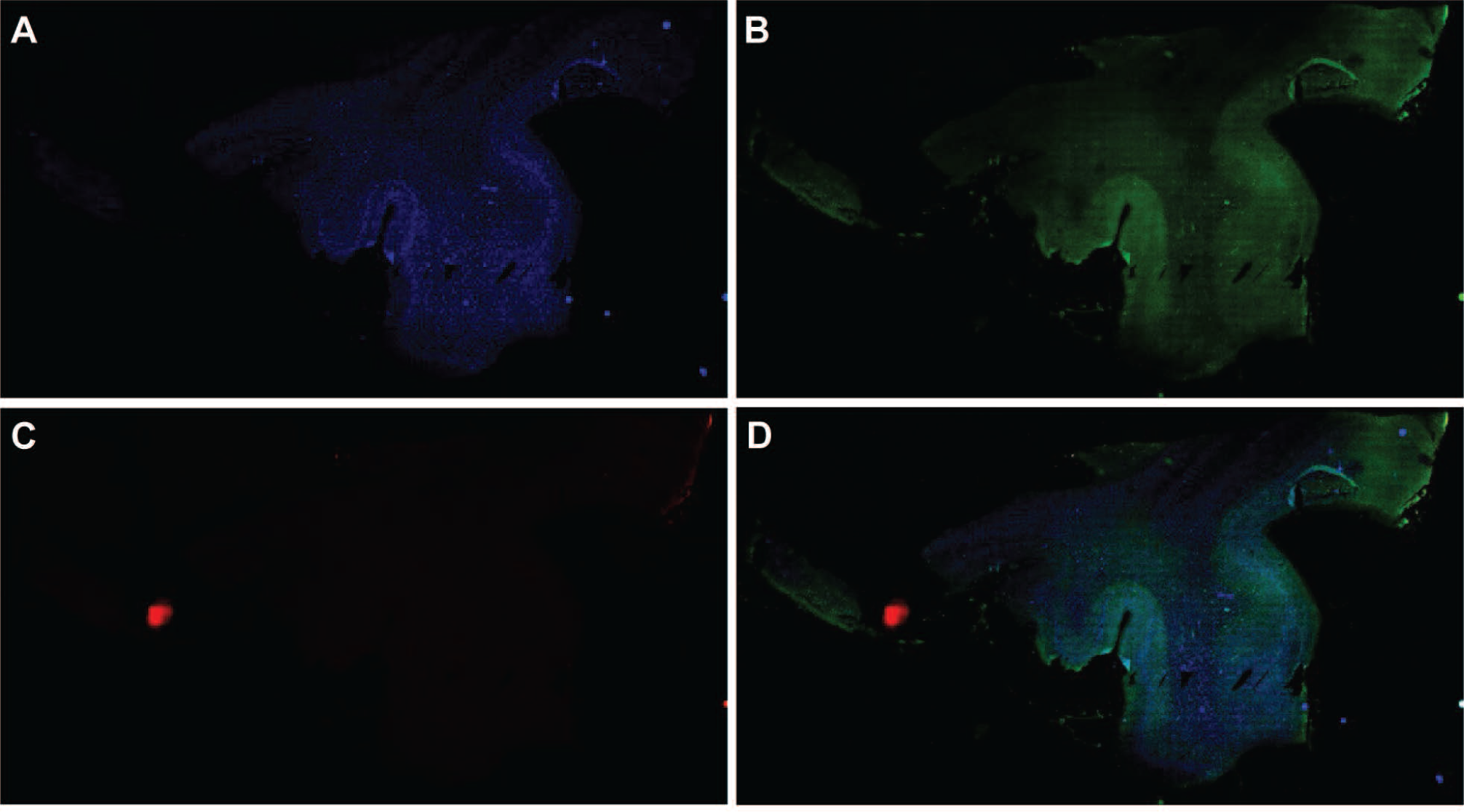
HSPA5 is more prevalent in the grey matter of the cortex from an Alzheimer’s disease donor. Single channel and overlay images from whole slide scan of a segment of cortex from an individual with Alzheimer’s disease Braak VI stained with DAPI, Proteintech 11587-1AP rabbit pAb HSPA5 (secondary 488) (secondary 488) and ProteinTech 66009-1-1 mouse mAb beta actin (secondary 568) antibodies. A: DAPI channel, B: 488 channel, C: 568 channel, D: overlay.

**Figure 11.**
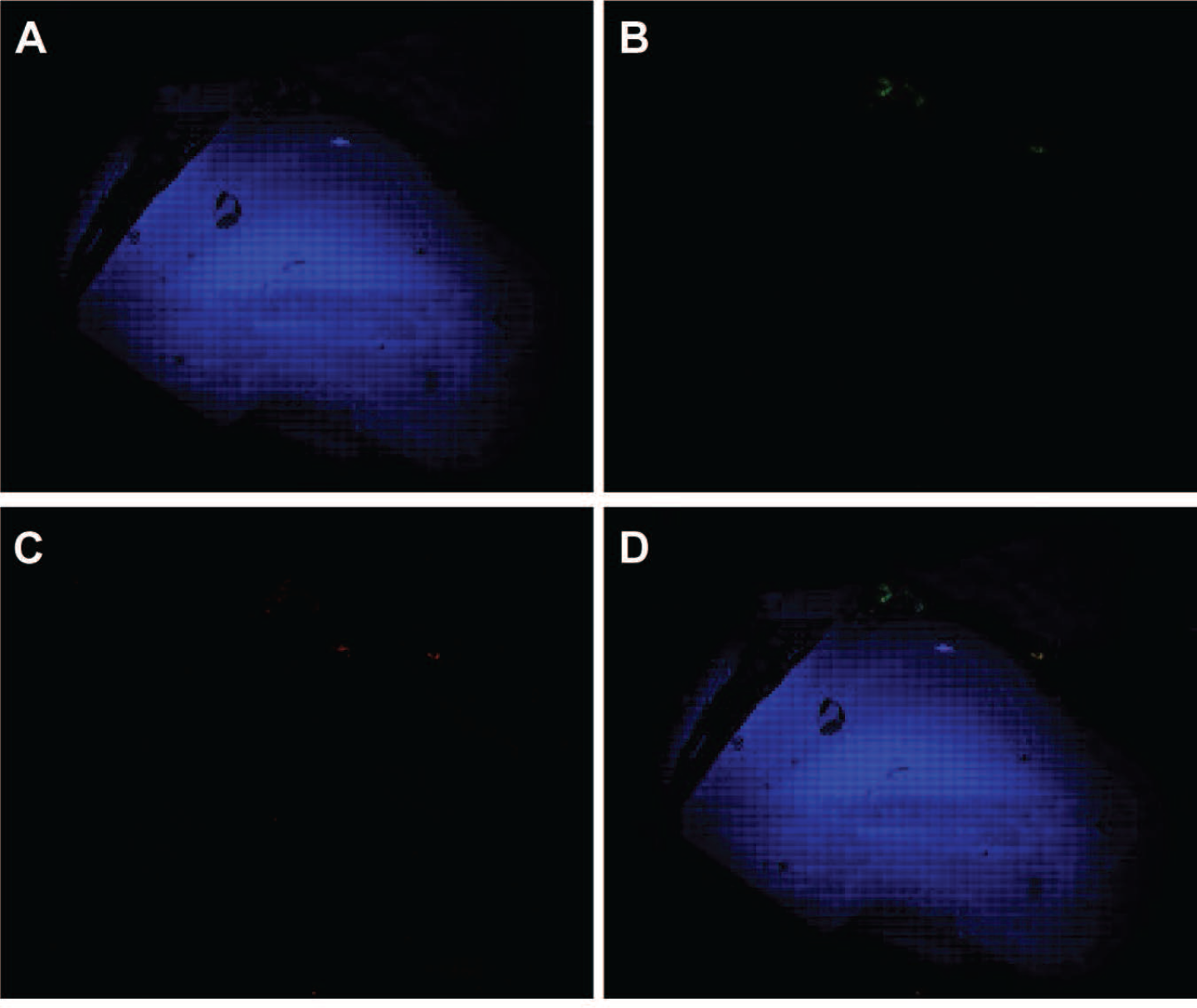
HSPA5 is more prevalent in the grey matter of the cortex from a Huntington’s disease donor. Single channel and overlay images from whole slide scan of a cortex FFPE sample from an individual with Huntington’s disease stained with DAPI, Proteintech 11587-1AP rabbit pAb HSPA5 (secondary 568), and Developmental Studies Hybridoma Bank MW7-s mouse mAb huntingtin (secondary 488) antibodies. A: DAPI channel, B: 488 channel, C: 568 channel, D: overlay.

**Figure 12.**
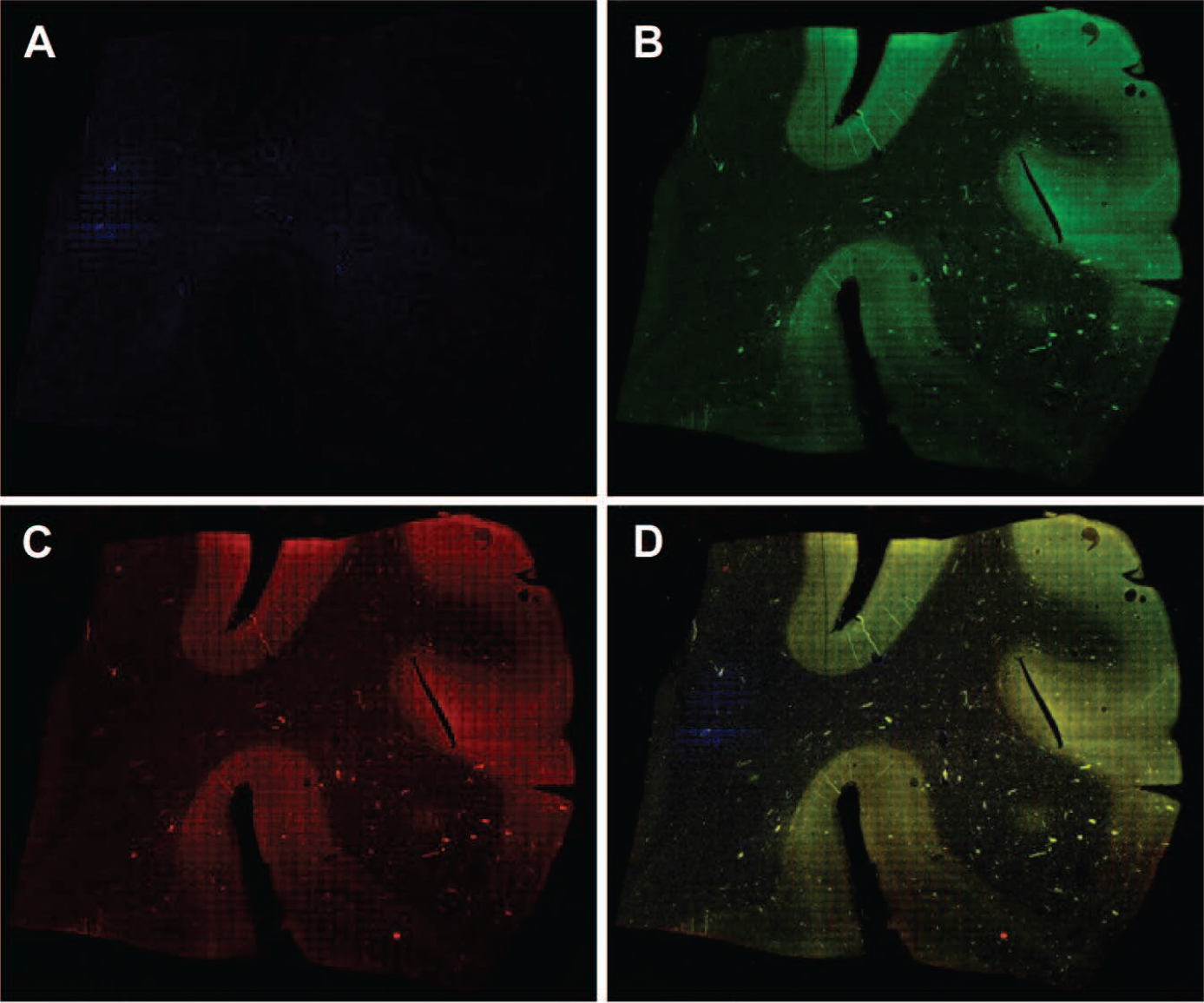
HSPA5 is more prevalent in the grey matter of the cortex from a Amyotrophic Lateral Sclerosis donor. Single channel and overlay images from whole slide scan of a cortex FFPE sample from an individual with Amyotrophic Lateral Sclerosis stained with DAPI, Proteintech 11587-1AP rabbit pAb HSPA5 (secondary 568) and Santa Cruz sc-515368 mouse mAb FICD clone G-7 (secondary 488) antibodies. A. DAPI channel, B: 488 channel, C: 568 channel, D: overlay.

**Figure 13.**
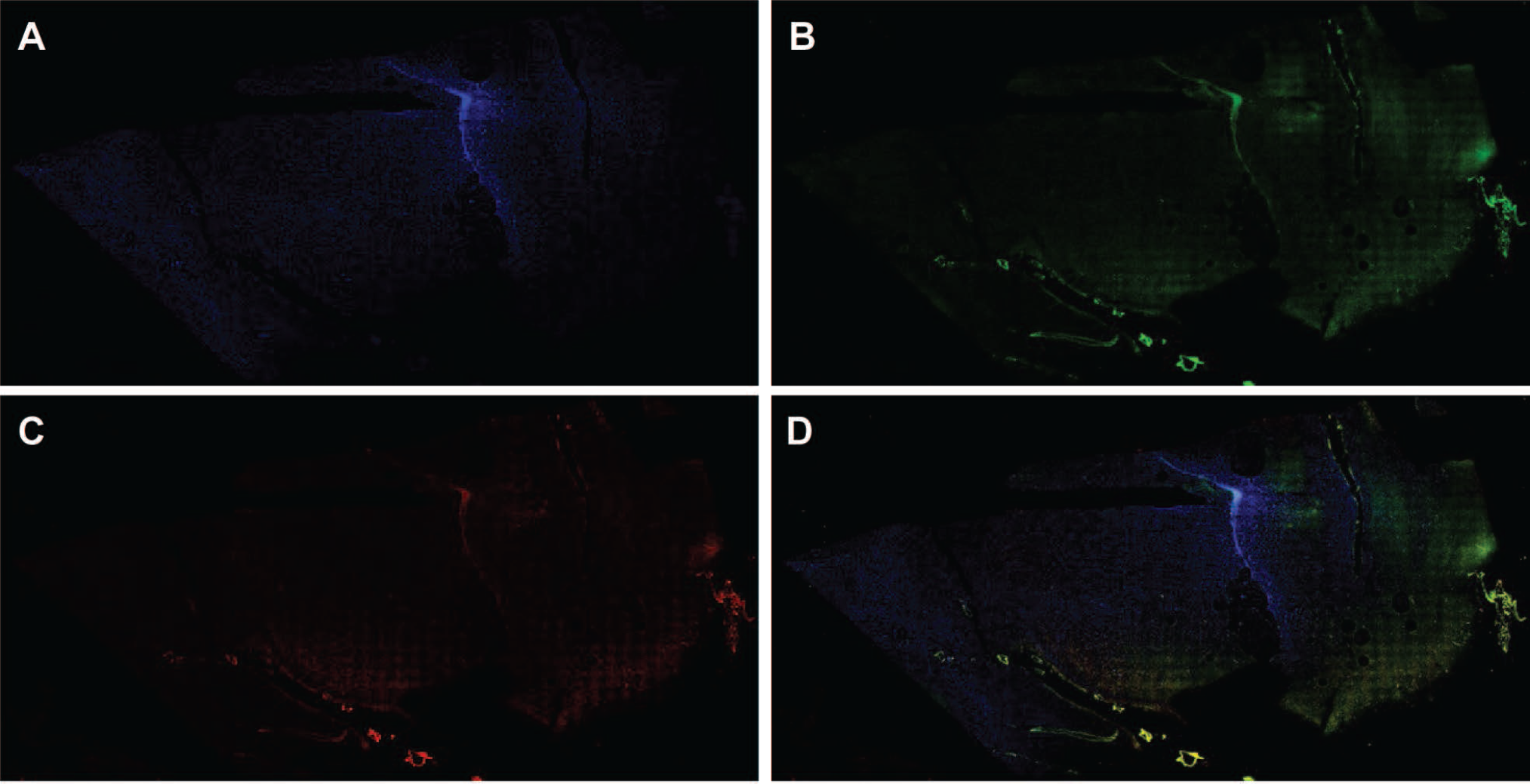
HSPA5 is more prevalent in the grey matter of the cortex from a donor with no neurological diagnosis. Single channel and overlay images from whole slide scan of a cortex FFPE sample from an individual with no neurological diagnosis stained with DAPI, Proteintech 11587-1AP rabbit pAb HSPA5 (secondary 568) and Santa Cruz sc-515368 mouse mAb FICD clone G-7 (secondary 488) antibodies. A: DAPI channel, B: 488 channel, C: 568 channel, D: overlay.

**Figure 14.**
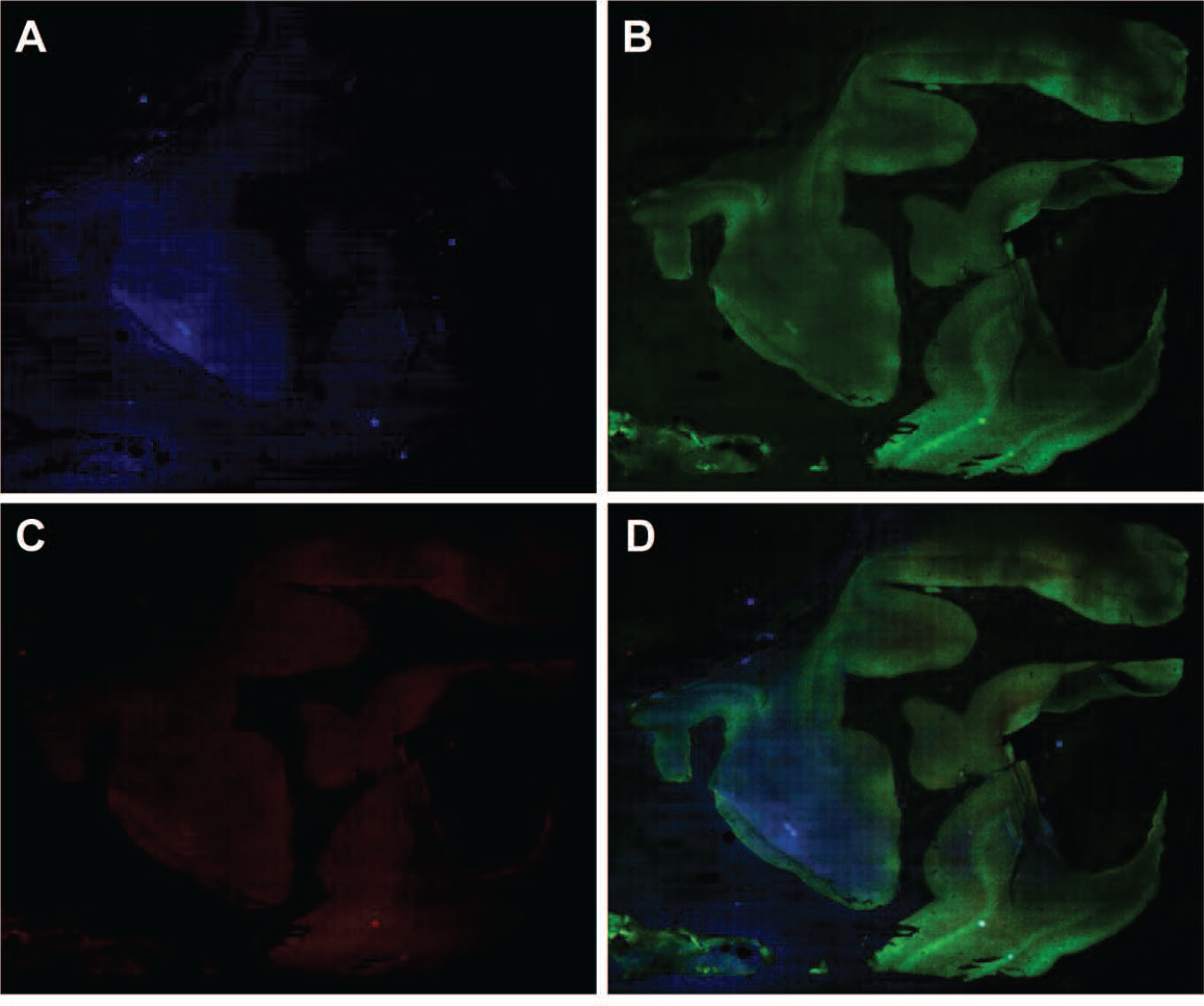
HSPA8 is abundant in the hippocampus in AD. HSPA8 is present in cells throughout a hippocampus from an Alzheimer’s disease donor, but these cells have a slightly higher concentration in the dentate gyrus. Single channel and overlay images from whole slide scan of a segment of hippocampus from an individual with Alzheimer’s disease Braak V/VI stained with DAPI, Santa Cruz 7298 mouse mAb HSPA8 (secondary 568) and Dako A0024 rabbit pAb Pan human tau (multiple isoforms) (secondary 488)antibodies. A: DAPI channel, B: 488 channel, C: 568 channel, D: overlay.

**Figure 15.**
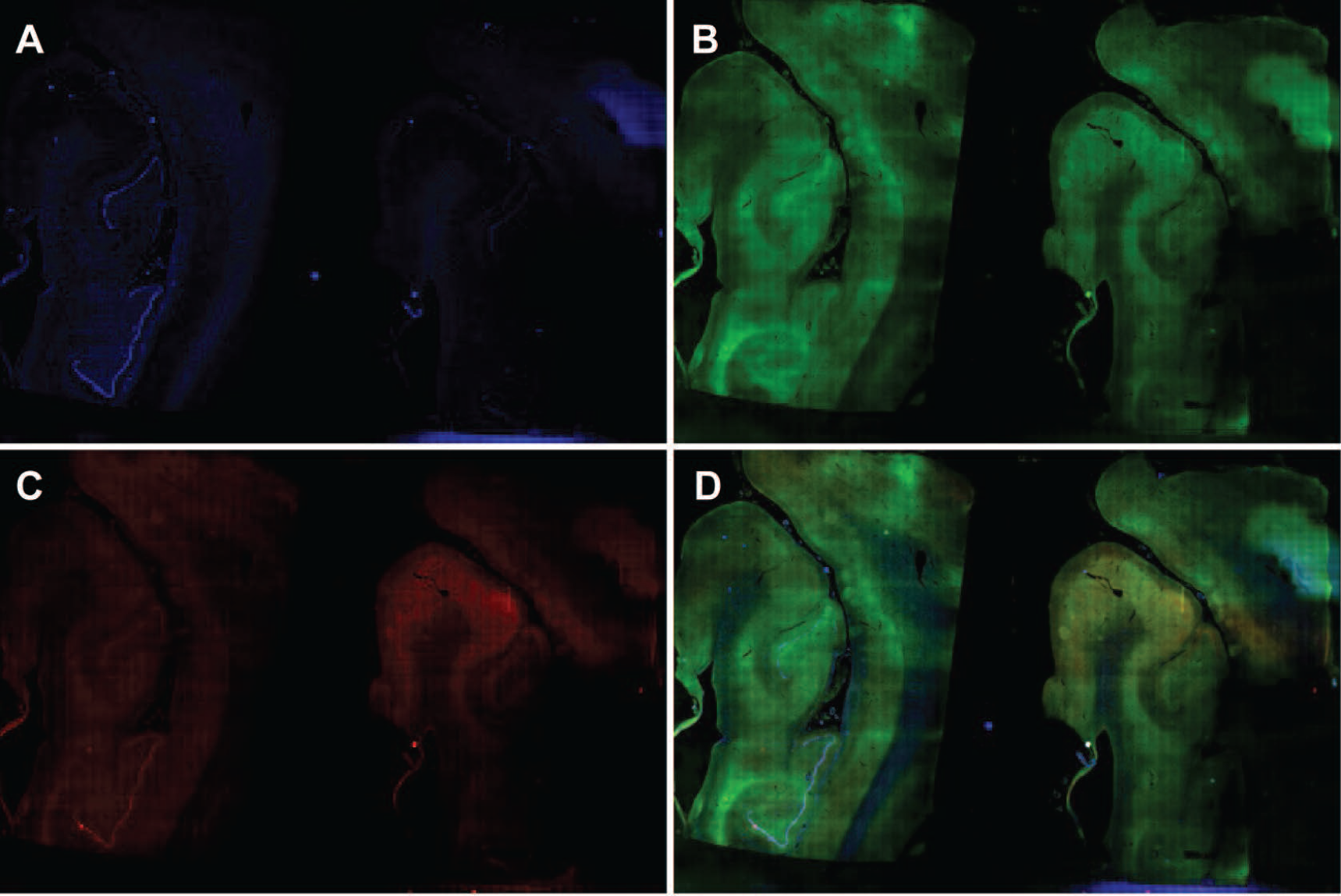
HSPA8 is abundant in the hippocampus in HD. HSPA8 is present in cells throughout a hippocampus from a Huntington’s disease donor, but these cells are more concentrated in the dentate gyrus. Single channel and overlay images from whole slide scan of a segment of hippocampus from an individual with Huntington’s disease stained with DAPI, Santa Cruz 7298 mouse mAb HSPA8 (secondary 568) and Dako A0024 rabbit pAb Pan human tau (multiple isoforms) (secondary 488) antibodies. A: DAPI channel, B: 488 channel, C: 568 channel, D: overlay.

**Figure 16.**
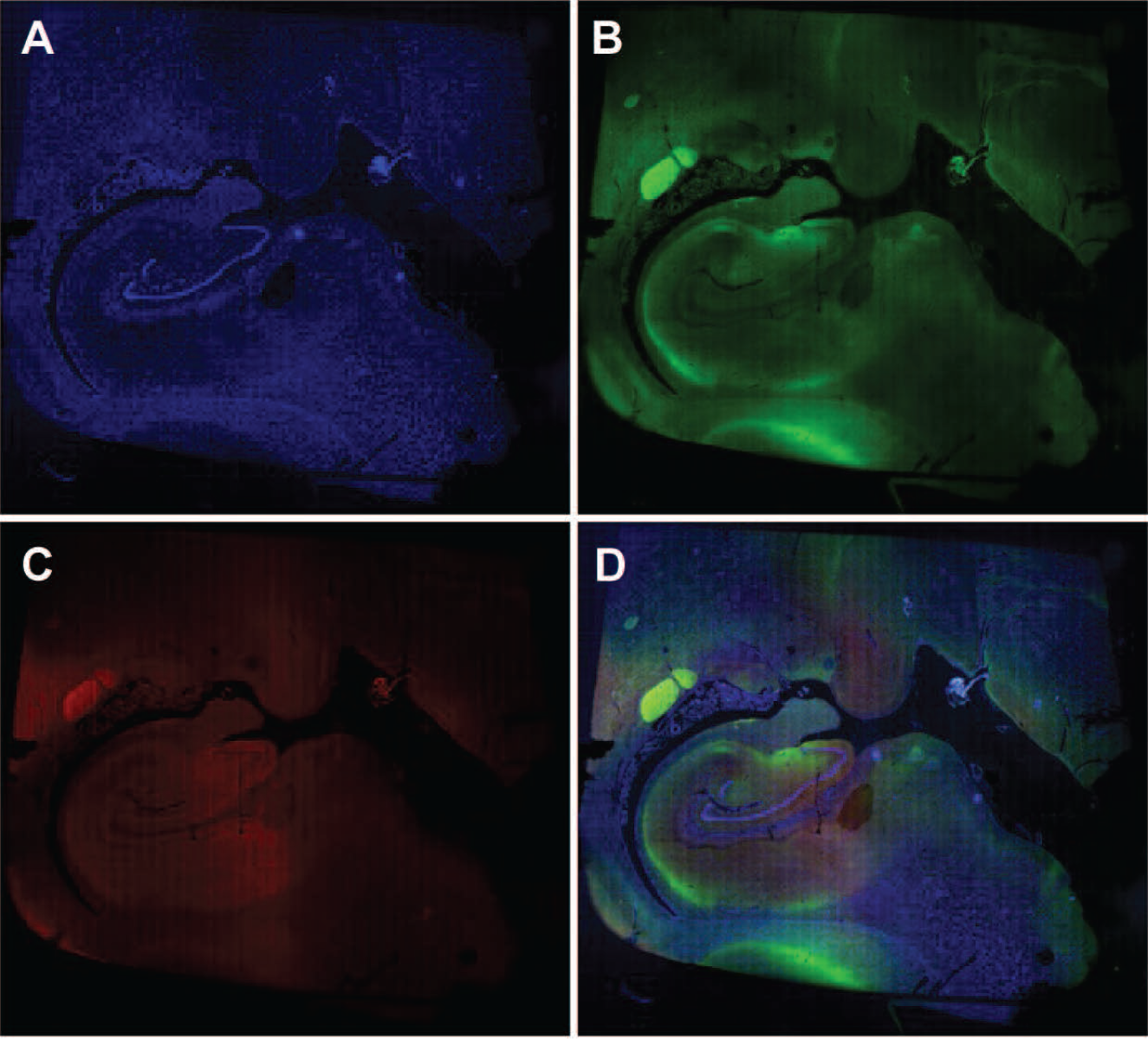
HSPA8 is abundant in the hippocampus in ALS. HSPA8 is present in cells throughout a hippocampus from an Amyotrophic Lateral Sclerosis donor, but these cells are more concentrated in the dentate gyrus. Single channel and overlay images from whole slide scan of a segment of hippocampus from an individual with Amyotrophic Lateral Sclerosis stained with DAPI, Santa Cruz 7298 mouse mAb HSPA8 (secondary 568) and Dako A0024 rabbit pAb Pan human tau (multiple isoforms) (secondary 488) antibodies. A: DAPI channel, B: 488 channel, C: 568 channel, D: overlay.

**Figure 17.**
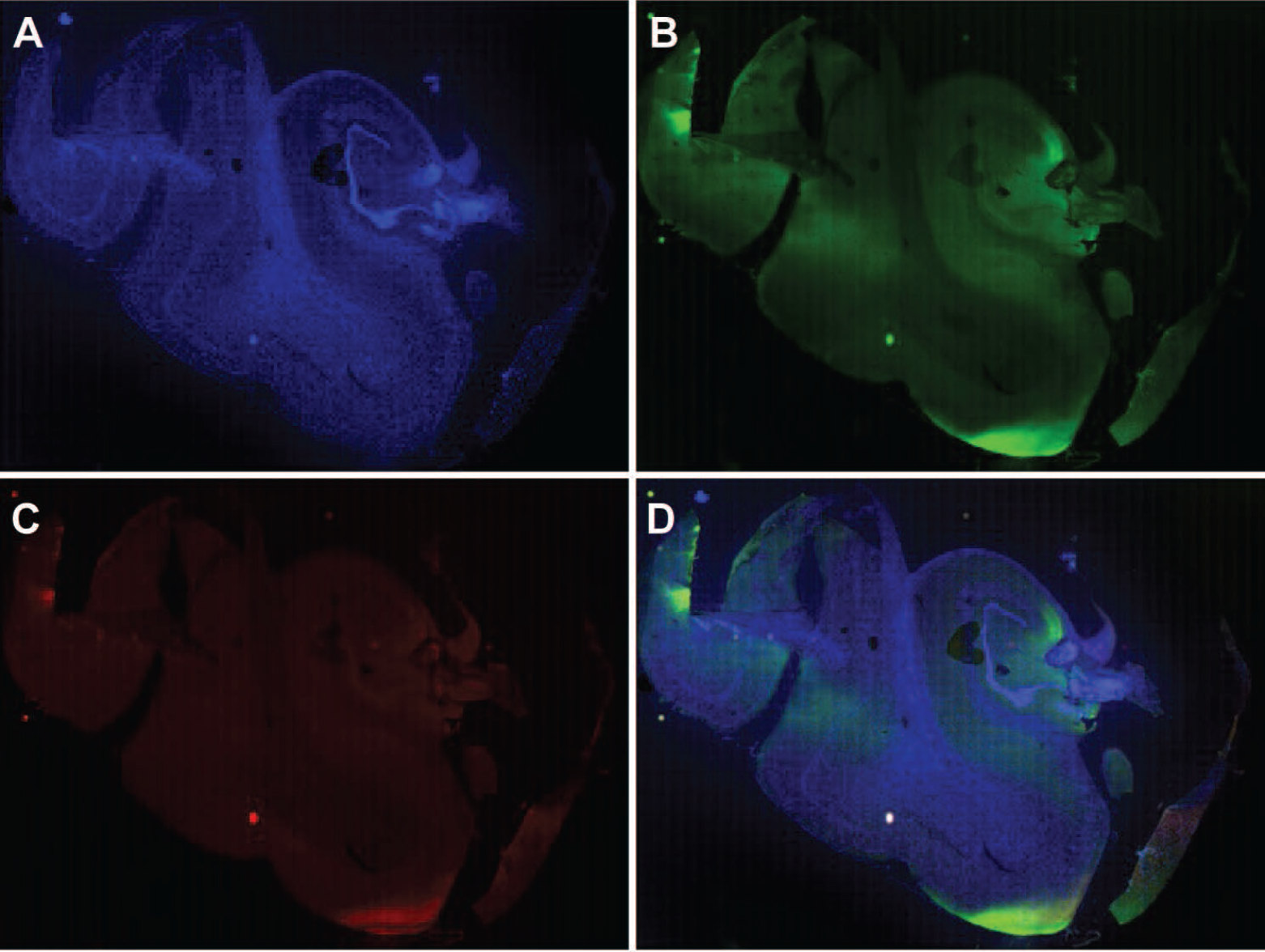
HSPA8 is abundant in the hippocampus from a donor with no neurological diagnosis. HSPA8 is present in cells throughout a hippocampus from a donor with no neurological diagnosis, but these cells are more concentrated in the dentate gyrus. Single channel and overlay images from whole slide scan of a segment of hippocampus from an individual with no neurological diagnosis stained with DAPI, Santa Cruz 7298 mouse mAb HSPA8 (secondary 568) and Dako A0024 rabbit pAb Pan human tau (multiple isoforms) (secondary 488) antibodies. A: DAPI channel, B: 488 channel, C: 568 channel, D: overlay.

**Figure 18.**
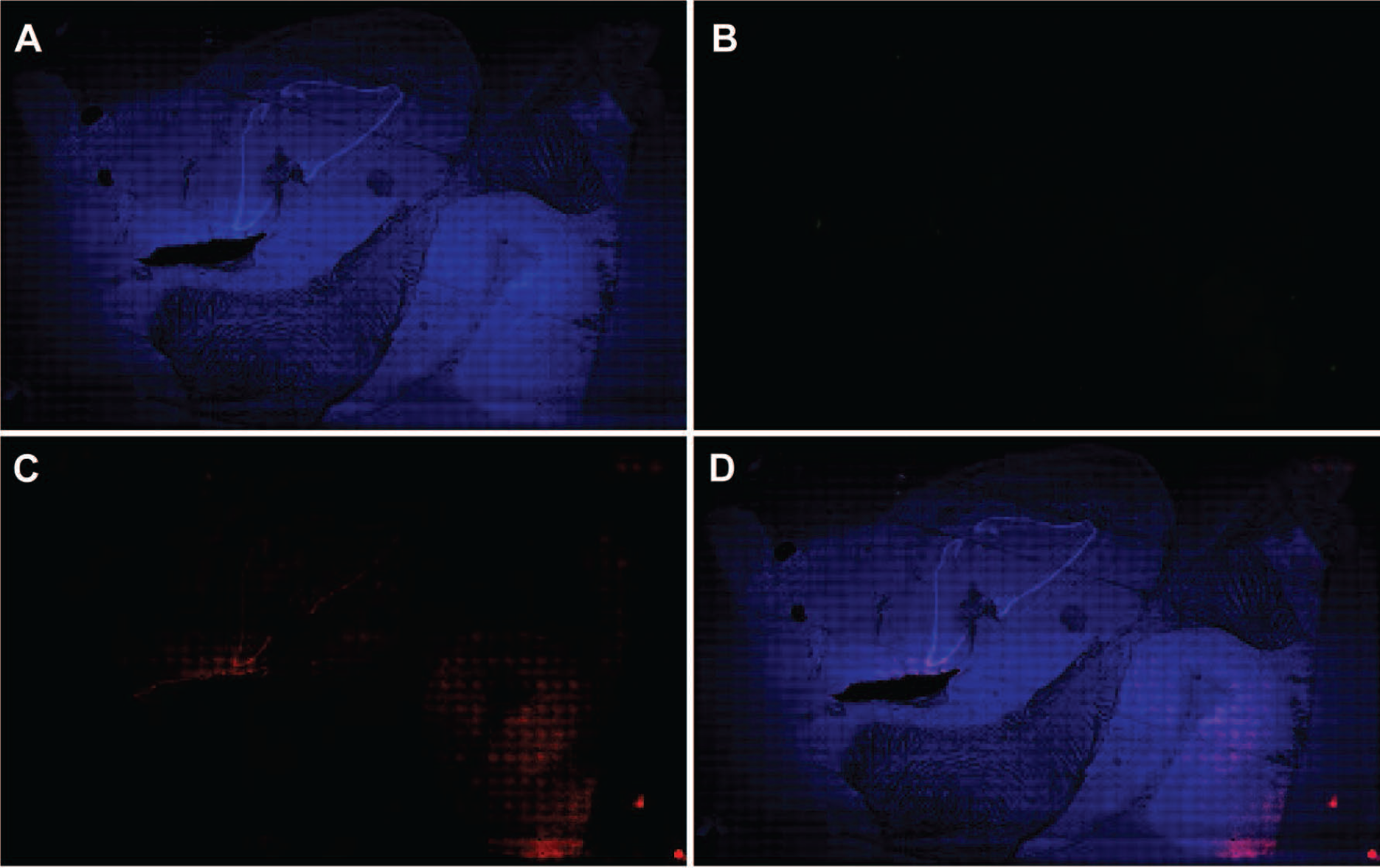
HSPA5 is abundant in the hippocampus in HD. HSPA5 is present in cells throughout a hippocampus from a Huntington’s disease donor, but these cells are more concentrated in the dentate gyrus. Single channel and overlay images from whole slide scan of a hippocampus from an individual with Huntington’s disease stained with DAPI, Proteintech 11587-1AP rabbit pAb HSPA5 (secondary 568) and Developmental Studies Hybridoma Bank MW7-s mouse mAb huntingtin at (secondary 488) antibodies. A: DAPI channel, B: 488 channel, C: 568 channel, D: overlay.

### Iterative staining allows for co-localization analysis of multiple targets

The main advantage of iterative staining is that it allows for the detection of more targets on the same piece of tissue compared to standard immunohistochemistry or immunofluorescence. These multiple images can then be used to determine how the various targets correspond to one another in the same sample. We compared images from subsequent staining cycles from an AD Braak IV/V hippocampus stained with anti-Neu antibody (secondary fluorophore 568) and anti-FICD antibody (secondary fluorophore 488) in round 1 and anti-Tau (multiple isoforms) antibody (secondary antibody 488) and anti-HSPA8 antibody (secondary fluorophore 568) used in cycle 2. In these images, we observed limited colocalization of NeuN and Tau (figure 19). This may be because Tau is normally expressed in the soma, whereas NeuN is located near the nucleus (*10, 11*). Additionally, in dying neurons, where Tau is more likely to be detected in the soma, NeuN expression is diminished (*10, 11*). FICD is sparsely expressed in this area, and HSPA8 is not expressed in this area of the hippocampus (figure 19).

**Figure 19.**
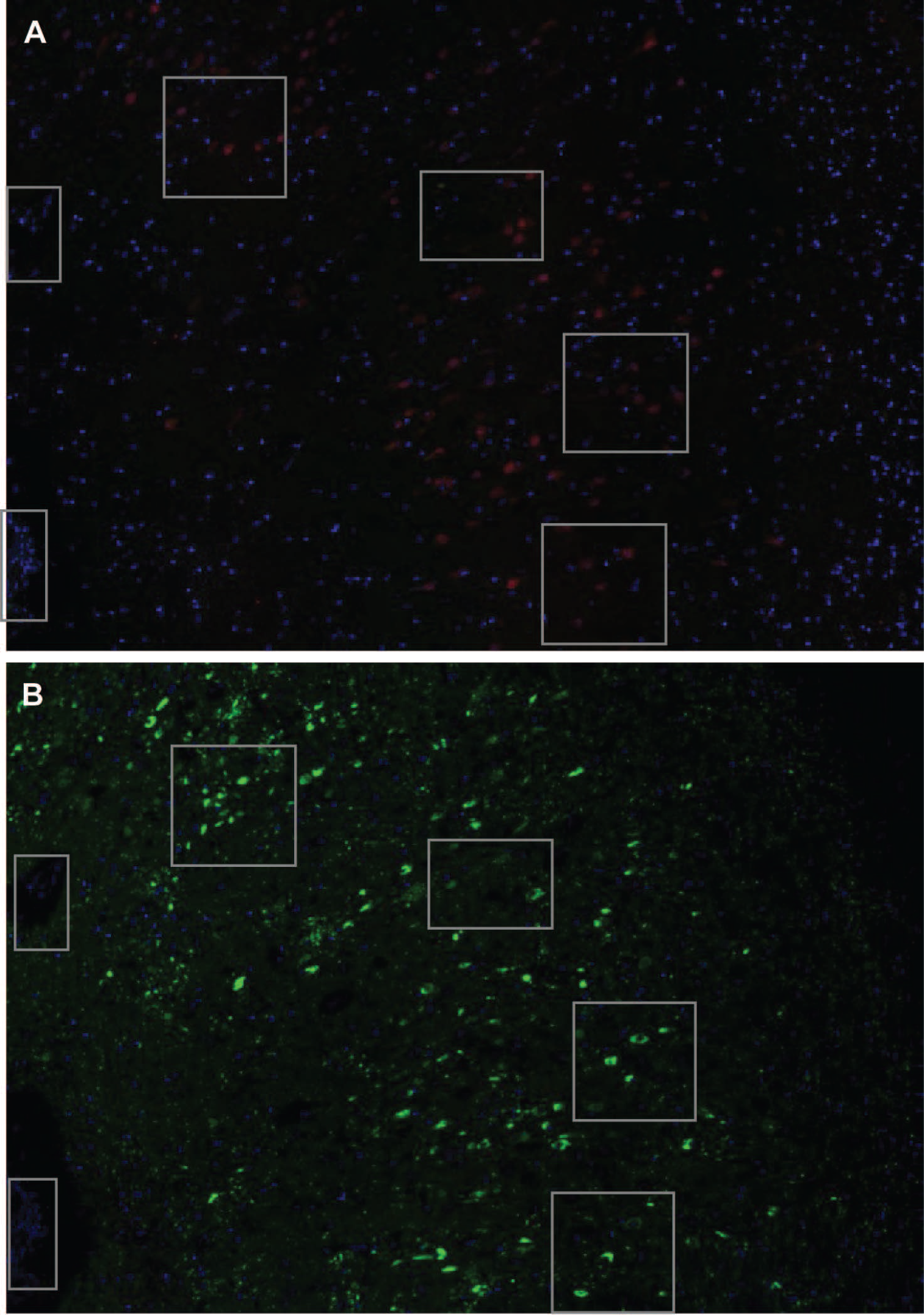
Tau only partially co-localizes with neuronal markers. Comparing cycles of staining shows that NeuN (neuronal nuclear marker) and Tau multiple isoforms (present in axons of neurons, astrocytes, and oligodendrocytes) co-localize to a small degree, with Tau being more evenly distributed in an area of a hippocampus near the dentate gyrus from a patient with Braak V/VI AD than NeuN. FICD is sparsely expressed in this area, and HSPA8 is not expressed in this area of the hippocampus. A section of hippocampus from an individual with Alzheimer’s Disease Braak V/VI was iteratively stained using the SHIELD protocol. Subsequent cycles are shown, with boxes highlighting corresponding areas. All cycles were stained with DAPI. A: Cycle 1 ProteinTech 26975-1-AP rabbit pAb NeuN (secondary 568) and Santa Cruz sc-515368 mouse mAb FICD clone G-7 (secondary 488) B: Cycle 2 DAKO A0024 Rabbit pAb Pan human tau (multiple isoforms) (secondary 488) Santa Cruz 7298 mouse mAb HSPA8 (secondary 568).

**Figure 20.**
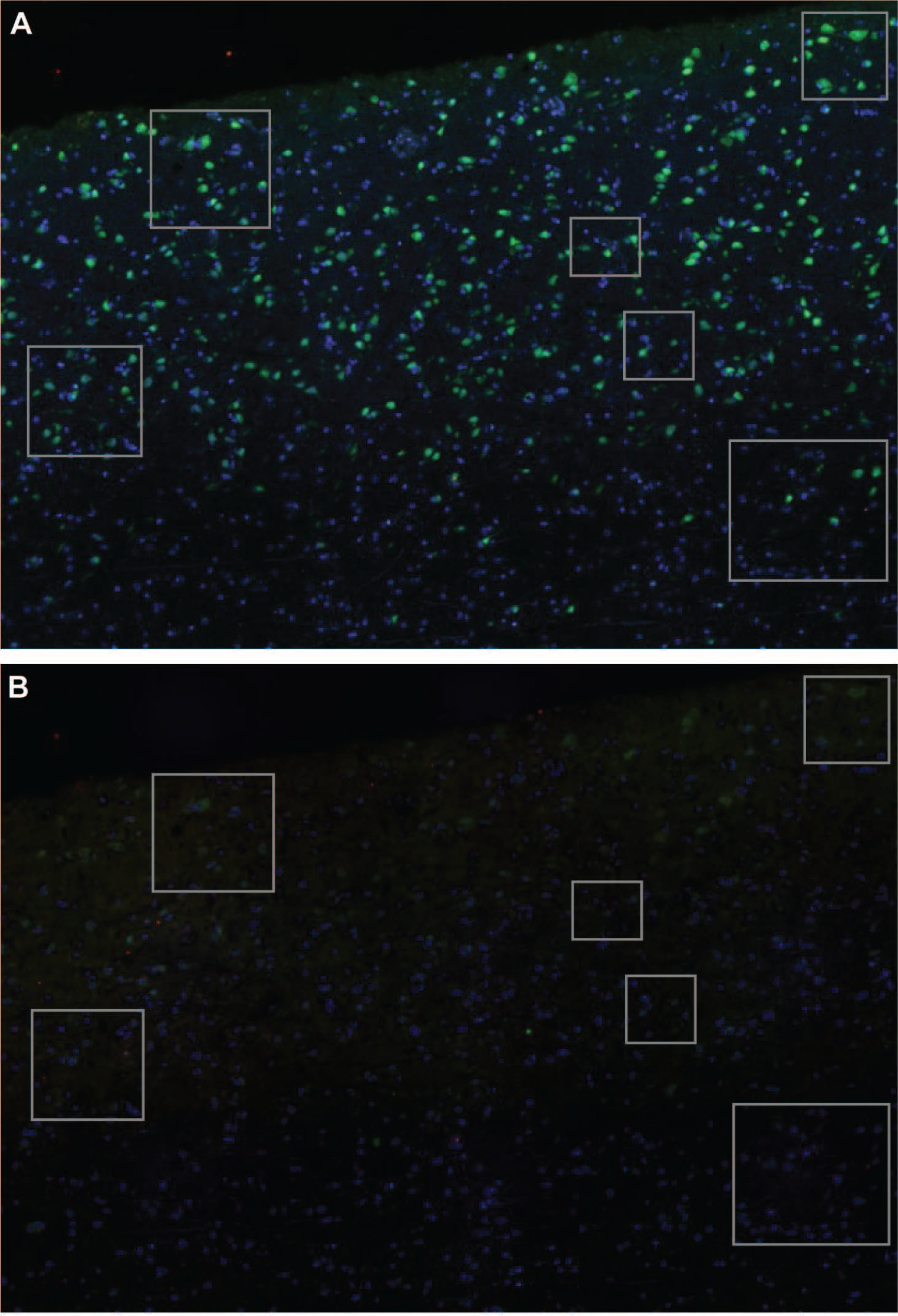
Tau only partially co-localizes with neuronal markers. Comparing cycles of staining shows that NeuN (neuronal nuclear marker) and Tau multiple isoforms (present in axons of neurons, astrocytes, and oligodendrocytes) co-localize to a moderate degree in an area of cortex from an individual with no neurological diagnosis. NeuN is more widely expressed in this region, but is more concentrated in the grey matter, as is Tau. Acetylcholinesterase (ACHE, cholinergic neuron marker) and HSPA8 are less abundant but are more concentrated in the grey matter as well. A section of cortex from an individual with no neurological diagnosis was iteratively stained using the SHIELD protocol. Subsequent cycles are shown, with boxes highlighting corresponding areas. All cycles were stained with DAPI. A: Cycle 2 ProteinTech 26975-1-AP rabbit pAb NeuN (secondary 488) and Developmental Studies Hybridoma Bank tor23 mouse anti-Acetylcholinesterase, presynaptic (secondary 568). B: Cycle 4 DAKO A0024 Rabbit pAb Pan human tau (multiple isoforms) (secondary 488) Santa Cruz 7298 mouse mAb HSPA8 (secondary 568).

## Discussion

In this study, we present a novel method, termed TSWIFT, to iteratively stain and image mounted FFPE-embedded cryosections of mammalian tissue. Our method allows for the staining of more than four target proteins in a single section, which is an often-found limit for traditional immunostaining methods using commercial primary and secondary antibodies. Next, TSWIFT can be performed using standard laboratory equipment. The ability to store samples for months between staining cycles offers the benefit to revisit previously imaged tissue samples and assess additional targets as they are identified during a study or as part of a revision process. We thus believe that our method will find broad application in distinct tissue-based research settings.

As expected, epoxidation strengthens the tissue, making repeat staining with minimal loss of tissue integrity possible. Gentle release of the coverslip further minimizes tissue disruption. However, the conservation of tissue integrity remains a key concern which warrants inspection prior to each staining cycle, as minor tissue tearing is likely to occur. Several software solutions for the location integration between staining cycles have already been developed by other groups, which maximizes the amount of information that can be gained from iterative staining approaches (*1, 2, 12, 13*). Most of these software solutions are specifically tailored to the staining technique used. We find that Correlia, developed by Rohde *et al.* (*12*), was the most adaptable to our technique. Based on our experience, the best results with localization integration are achieved if the input images were acquired on a confocal microscope, which may represent an additional constrain to some laboratories. Despite some limitations, this technique has great potential.

Using TSWIFT, we examined chaperone localization across *post mortem* human cortex and cerebellum tissue from neurologically unaffected controls as well as ALS, Alzheimer’s disease and Huntington’s disease patients. We find that HSPA8, and to a lesser extend HSPA5, are both enriched in grey matter (cortex) and dentate nucleus (cerebellum). HSPA8 is the cognate Hsp70 family chaperone (HSC70), required for *de novo* protein folding, protein recycling, chaperone-mediated autophagy, protein disaggregation, proteosomal protein degradation. HSPA5 is the ER resident HSP70 family chaperone (BiP) which is essential for the proper folding of secreted proteins that pass through the ER. Both chaperones were proposed to be expressed in high levels in all cells. While our study was not sufficiently powered to identify and establish disease-specific changes in chaperone abundance and localization, we find that that HSPA8 and HSPA5 levels vary across brain tissues. A plausible interpretation of this result could be that HSPA8 and HSPA5 abundance varies across cell types, with secretory cells containing increased levels. Further investigation will be needed to address these questions in sufficient detail.

## Acknowledgments

We thank the members of the Truttmann lab for helpful comments and discussion. William Giblin is acknowledged for proof-reading the manuscript draft. Shannon Lacy, Nicholas Urban, and William Giblin are acknowledged for defining the method’s name (TSWIFT). CMP was supported by NIA Training Grant AG000114. MCT is supported by an Alzheimer’s foundation Young Investigator Award, a Ruth K Broad foundation award and grant 1R35GM142561.

## Author contributions

MCT supervised the project. MCT and CMP planned and designed the experiments. CMP performed all experiments. CMP and MCT wrote the manuscript.

## Data availability statement

The unprocessed raw datasets (images) generated and analyzed during the current study are available from the corresponding author upon reasonable request.

## Additional information

The authors declare no competing interests.

---

**Supplementary Figure S1. Sudan Black is necessary to reduce background fluorescence.** A: Whole slide scan of cycle 2 of staining on a section of cortex from an individual with no neurological diagnosis stained with DAPI, ProteinTech 26975-1-AP rabbit pAb NeuN (secondary 488) and Developmental Studies Hybridoma Bank tor23 mouse mAb Acetylcholinesterase, presynaptic (secondary 568) using Sudan Black. B: The same section on cycle 3 stained with no primaries or secondaries with no Sudan Black.

**Supplementary Figures S2-8: SHIELD protects tissue and allows for repeated staining of tissue with maintained signal.** 20x images from cycles of staining and destaining post-SHIELD. All are stained with DAPI, Santa Cruz sc-58860 mouse mAb Tau (secondary 568) and DAKO Z0334 rabbit pAb GFAP (secondary 488). Unless otherwise noted, images were taken at the exposure automatically selected by the BZ-X700. Because of this, images of destains were taken at much higher exposure than those of stains to detect low amounts of signal. Fig S2: cycles 1 and 2 stained and destained. Fig S3: cycles 3 and 4 stained and destained. Fig S4: cycles 5 and 6 (5 is secondaries only and was taken at the exposure settings used for cycle 4, images were not taken after the cycle 5 destain). Fig S5: cycles 7 and 8. Fig S6: cycles 9 and 10 (in round 10, secondary antibody fluorophores were swapped, images were not taken after the cycle 10 destain). Fig S7: cycles 11 and 12 (no destain images were taken for cycle 12). Fig S8: cycles 13 and 14 (no destain images for either round).

