## Supplemental Table 1 for "TSWIFT, a novel method for iterative staining of embedded and mounted human brain sections"

| Target | Company | Raised in | Catalog Numbe | Concentration use |
|---|---|---|---|---|
| HSP27/HSPB1 | Cell Signaling | mouse | 2402 | 1:50 |
| Hsp60 | Cell Signaling | rabbit | 12165 | 1:500 |
| total HSP70 and HSPA8 | Cell Signaling | rabbit | 4872 | 1:50 |
| β-amyloid (pE3 peptide) | Cell Signaling | rabbit | 14975 | 1:100 |
| Glial Fibrillary Acid Proteir | Dako | rabbit | Z0334 | 1:500 |
| Pan human tau | Dako | rabbit | A0024 | 1:1,000 |
| ACHE acetylcholinesteras | Developmental Studies Hybridoma Ban | mouse | tor23 | 5ug/mL |
| alpha-tubulin | Developmental Studies Hybridoma Ban | mouse | 12G10 | 5ug/mL |
| HTT | Developmental Studies Hybridoma Ban | mouse | MW-7 | 5ug/mL |
| HTT aggregate | Developmental Studies Hybridoma Ban | mouse | MW-8 | 5ug/mL |
| Thr-AMP | Lampire | rabbit | custom-made | 1:50 |
| ATF-6B | Proteintech | rabbit | 15794-1-AP | 1:100 |
| Beta actin | Proteintech | mouse | 66009-1-1 | 1:1,000 and 1:50 |
| HSPA5/BiP/GRP78 | Proteintech | rabbit | 11587-1AP | 1:100 |
| NeuN | Proteintech | rabbit | 26975-1 | 1:1,000 |
| alpha-synuclein | Santa Cruz Biotechnology | mouse | sc-12767 | 1:50 |
| CD68 | Santa Cruz Biotechnology | mouse | sc-17832 | 1:50 |
| DARPP-32 | Santa Cruz Biotechnology | mouse | sc-271111 | 1:50 |
| FICD/HYPE | Santa Cruz Biotechnology | mouse | sc-515368 | 1:50 |
| Flk-1/KDR/VEGFR2 | Santa Cruz Biotechnology | mouse | sc-6251 | 1:50 |
| HSPA8/Hsc70 | Santa Cruz Biotechnology | mouse | sc-7298 | 1:50 |
| pan Tau | Santa Cruz Biotechnology | mouse | sc-58860 | 1:100 |
