## Supplementary figures and images for "TSWIFT, a novel method for iterative staining of embedded and mounted human brain sections"

### Supplemental figures S1-S3

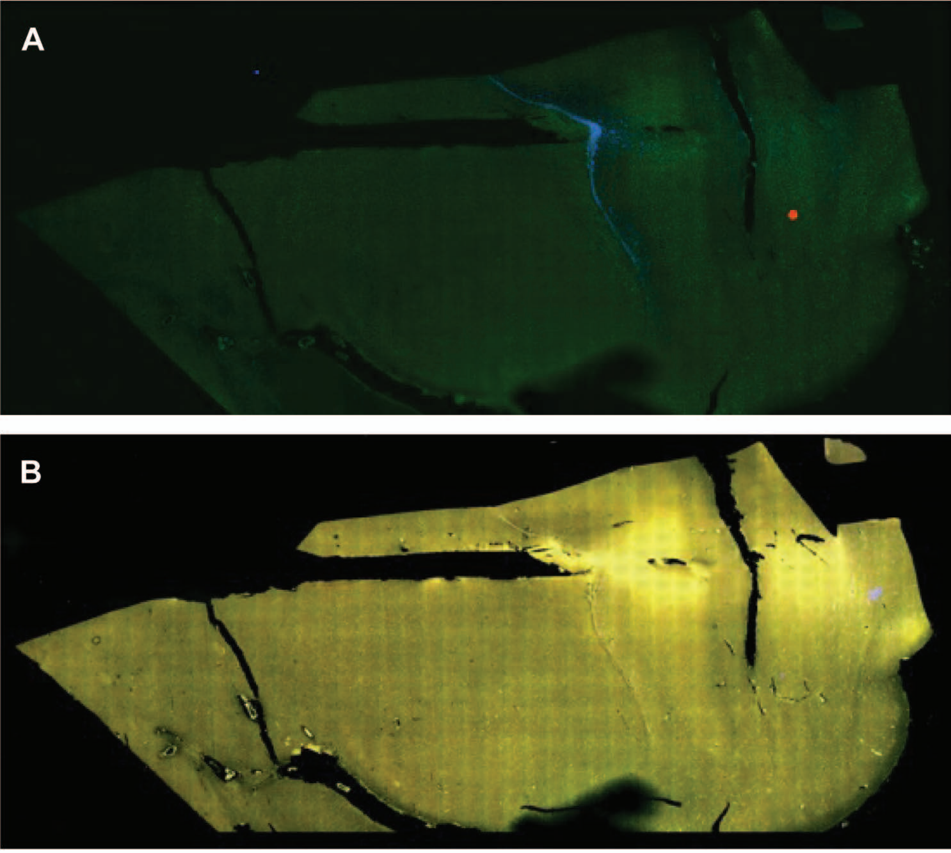


**Figure S1**

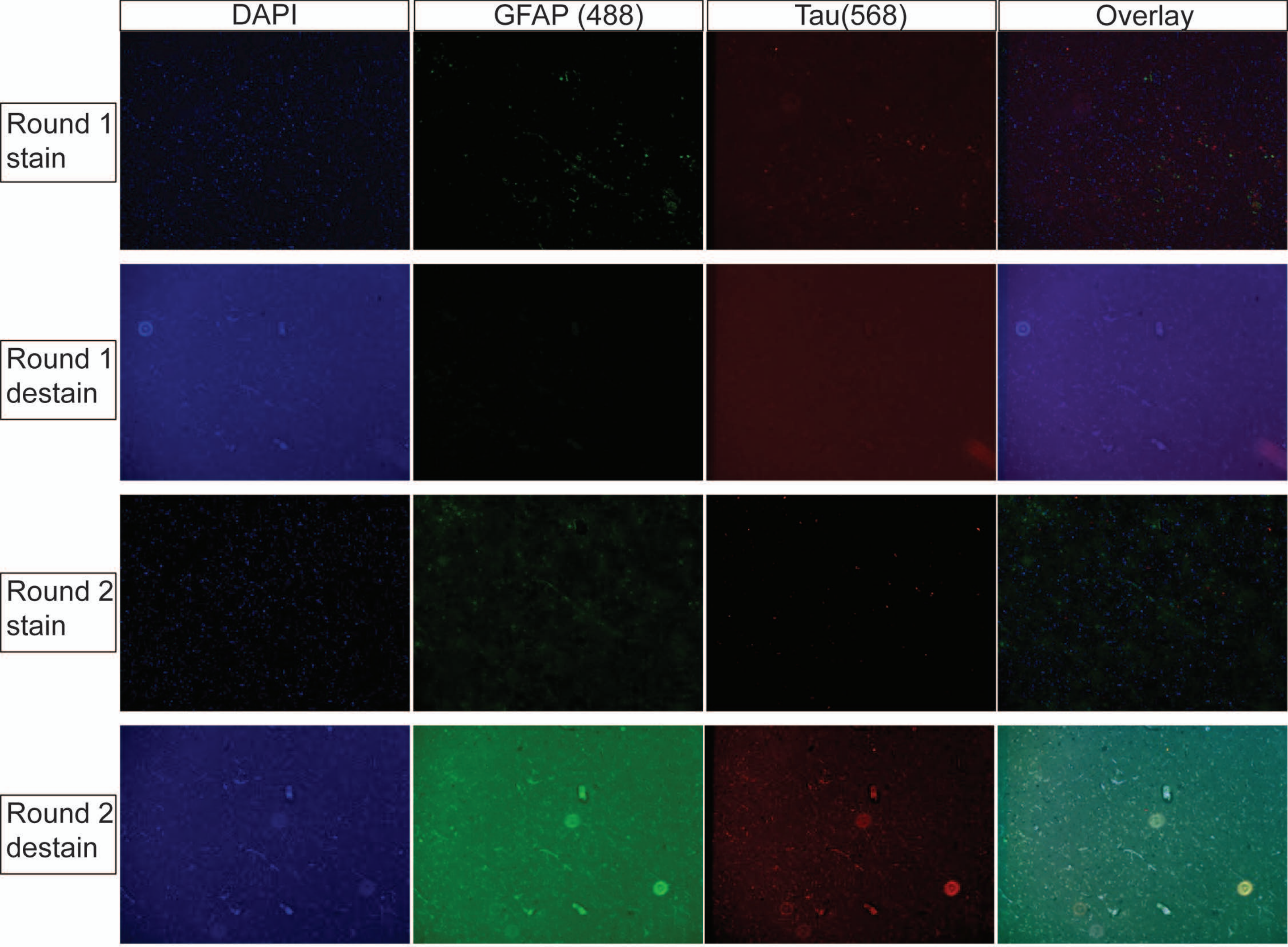

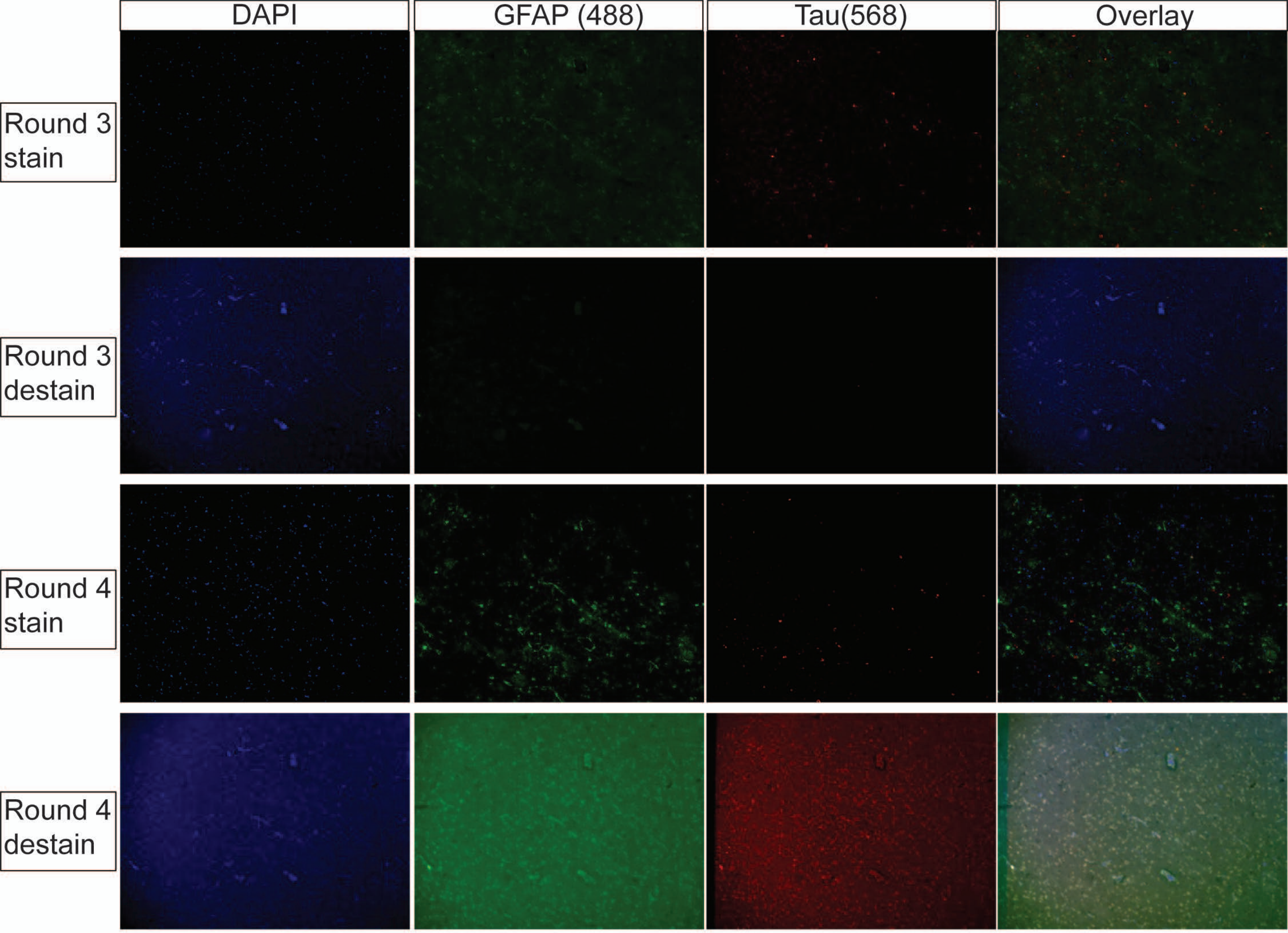
